# Predicting environmental stressor levels with machine learning: a comparison between amplicon sequencing, metagenomics, and total RNA sequencing based on taxonomically assigned data

**DOI:** 10.1101/2022.11.18.517107

**Authors:** Christopher A. Hempel, Dominik Buchner, Leoni Mack, Marie V. Brasseur, Dan Tulpan, Florian Leese, Dirk Steinke

## Abstract

**Background:** Microbes are increasingly (re)considered for environmental assessments because they are powerful indicators for the health of ecosystems. The complexity of microbial communities necessitates powerful novel tools to derive conclusions for environmental decision-makers, and machine learning is a promising option in that context. While amplicon sequencing is typically applied to assess microbial communities, metagenomics and total RNA sequencing (herein summarized as omics-based methods) can provide a more holistic picture of microbial biodiversity at sufficient sequencing depths. Despite this advantage, amplicon sequencing and omics-based methods have not yet been compared for taxonomy-based environmental assessments with machine learning. In this study, we applied 16S and ITS-2 sequencing, metagenomics, and total RNA sequencing to samples from a stream mesocosm experiment that investigated the impacts of two aquatic stressors, insecticide and increased fine sediment deposition, on stream biodiversity. We processed the data using similarity clustering and denoising (only applicable to amplicon sequencing) as well as multiple taxonomic levels, data types, feature selection, and machine learning algorithms and evaluated the stressor prediction performance of each generated model for a total of 1,536 evaluated combinations of taxonomic datasets and data-processing methods.

**Results:** Sequencing and data-processing methods had a substantial impact on stressor prediction. While omics-based methods detected much more taxa than amplicon sequencing, 16S sequencing outperformed all other sequencing methods in terms of stressor prediction based on the Matthews Correlation Coefficient. However, even the highest observed performance for 16S sequencing was still only moderate. Omics-based methods performed poorly overall, but this was likely due to insufficient sequencing depth. Data types had no impact on performance while feature selection significantly improved performance for omics-based methods but not for amplicon sequencing.

**Conclusion:** Amplicon sequencing might be a better candidate for machine-learning-based environmental stressor prediction than omics-based methods, but the latter require further research at higher sequencing depths to confirm this conclusion. More sampling could improve stressor prediction performance, and while this was not possible in the context of our study, thousands of sampling sites are monitored for routine environmental assessments, providing an ideal framework to further refine the approach for possible implementation in environmental diagnostics.

## 1 Background

Globally, ecosystems are experiencing an unprecedented amount of human-induced environmental stress, in particular caused by climate change, land use, pollution, habitat fragmentation, and the introduction of invasive species. As a consequence, ecosystems are deteriorating and biodiversity is declining faster than ever before in human history [1–3]. The loss of biodiversity has extremely negative effects on ecosystem functions and, thereby, ecosystem services, which also reduces the economic value of ecosystems [4]. As a consequence, environmental management to protect and restore ecosystems has garnered increased attention, also at the political level [1].

Environmental management includes the identification of prevalent stressors and their impacts on ecosystem health. Very good indicators of ecosystem health are microbes (prokaryotes and unicellular eukaryotes) because they play a crucial role in ecosystems and are extremely sensitive to changes in environmental conditions. Consequently, their community composition can reveal important information about the health and stress levels of ecosystems, which can be utilized for routine biomonitoring to guide measures for the protection and restoration of ecosystems [5–8]. Microbial community composition is usually determined by using amplicon sequencing, which involves target PCR to amplify taxonomic barcode genes (amplicons), typically the 16S ribosomal RNA (rRNA) gene for prokaryotes, the internal transcribed spacer 2 (ITS-2) 2 for fungi, and the 18S rRNA gene for other microbial eukaryotes. Although this approach can introduce taxonomic and abundance bias due to varying binding affinities and amplification efficiencies of target primers [9– 14], amplicon sequencing is widely used because it is comparably cheap and can generate valuable and consistent information on community composition.

In contrast, metagenomics and metatranscriptomics are target-PCR-free methods that are usually applied to analyze the presence and expression of functional genes within communities [15– 18]; however, both methods also generate valuable data that can be used for taxonomic identification of community members as an alternative to amplicon sequencing.

Metagenomics targets all DNA in a sample, including non-functional genes, repetitive regions, and genes containing little taxonomic information due to insufficient variation. A vast number of these genes is lacking reference sequences in databases, and therefore, metagenomics generates large amounts of sequences that cannot be taxonomically annotated. At insufficient sequencing depth, this leads to a low biodiversity coverage that is outperformed by that of amplicon sequencing [9,19,20]. However, this limitation can be overcome by increasing the sequencing depth, and if the depth is increased sufficiently, biodiversity coverage through metagenomics can outperform that of amplicon sequencing [21–24].

Total RNA sequencing (total RNA-Seq) [25–27], also termed double-RNA approach [28], metatranscriptomics analysis of total rRNA [29], total RNA metatranscriptomics [30], or total RNA-seq-based metatranscriptomics [27], refers to metatranscriptomics without an mRNA enrichment step. Cellular RNA consists mostly of rRNA, including 16S and 18S rRNA, which means that a large portion of total RNA-Seq data can be used for taxonomic annotations of microbes. In a previous study, we showed that total RNA-Seq can identify a microbial mock community consisting of 10 species more accurately than metagenomics at almost one order of magnitude lower sequencing depth [31]. Therefore, total RNA-Seq combines the advantages of both amplicon sequencing and metagenomics, as it avoids targeted PCR while producing large amounts of 16S and 18S sequences that can be taxonomically annotated.

Both Metagenomics and metatranscriptomics are more costly than amplicon sequencing but they can deliver target-PCR-free functional and taxonomical information across the tree of life, and as a result, there is a growing interest in their application for ecological assessments [7,32–34].

Another area gaining increased interest for ecological assessments is machine learning. Machine learning comprises algorithms to discover structural patterns in data that can be used to make predictions. Learning, in that sense, means that the applied algorithms change their behavior through training so that they perform better going forward [35]. Machine learning is increasingly being applied in biological sciences, including microbial ecology and environmental assessments, due to its capacity to deal with the expanding scale and complexity of biological data [36,37]. Cordier et al. (2019) stated that machine learning is the most promising approach for routine biomonitoring as it has the potential to be faster, more cost-efficient, and more accurate than current morphology-based methods, and some researchers believe that ecology represents one of the most relevant areas for machine learning because it could solve a wide and diverse variety of ecological problems [38]. It already has been applied successfully to amplicon-sequencing-based environmental assessments in freshwater [5,39], marine and coastal water [40–45], and soil [46], overcoming both the complex biological challenges associated with environmental data and the statistical challenges associated with the interpretation of large datasets. However, for the prediction of ecological variables with taxonomically assigned metagenomic data, machine learning has been applied only once so far [47] and not at all with total RNA-Seq data. At this point, High-Throughput Sequencing (HTS) has reached sequencing depths that allow for the application of omics-based approaches in environmental studies; however, it is unclear what scale is required to allow for predictive machine learning approaches such as environmental stressor prediction. There is a clear need for a comparative assessment of metagenomics, total RNA-Seq, and amplicon sequencing with respect to their ability to provide adequate taxonomic datasets for machine learning approaches.

In this study, we compare the performance of amplicon sequencing, metagenomics, and total RNA-Seq to predict environmental stressor levels based on taxonomically assigned data using machine learning. We used samples obtained from an ExStream system [48] consisting of stream mesocosms that were exposed to fine sediment and an insecticide to investigate the impact of these aquatic key stressors on stream biodiversity and decomposition of organic [49] [note to reviewers: this citation is in submission and will be updated once it is published]. For amplicon sequencing, we used the two marker genes ITS-2 and 16S both with an operational taxonomic unit (OTU) clustering and an exact sequence variant (ESV) denoising method. We evaluated the markers individually as well as combined (multi-marker approach). Stressor prediction performance (SPP) for all datasets was based on different taxonomic levels (phylum, class, order, family, genus, and species), data types (abundance, presence–absence (P–A)), feature selection (with feature selection, without feature selection), and machine learning algorithms (k-Nearest Neighbors, Linear Support Vector Classification, Logistic Ridge Regression, Logistic Lasso Regression, Multilayer Perceptron, Random Forest, Support Vector Classification, and XGBoost).

## 2 Material and Methods

The overall study design is shown in Figure 1, and further details are given in the balance of this section.

**Figure 1:**
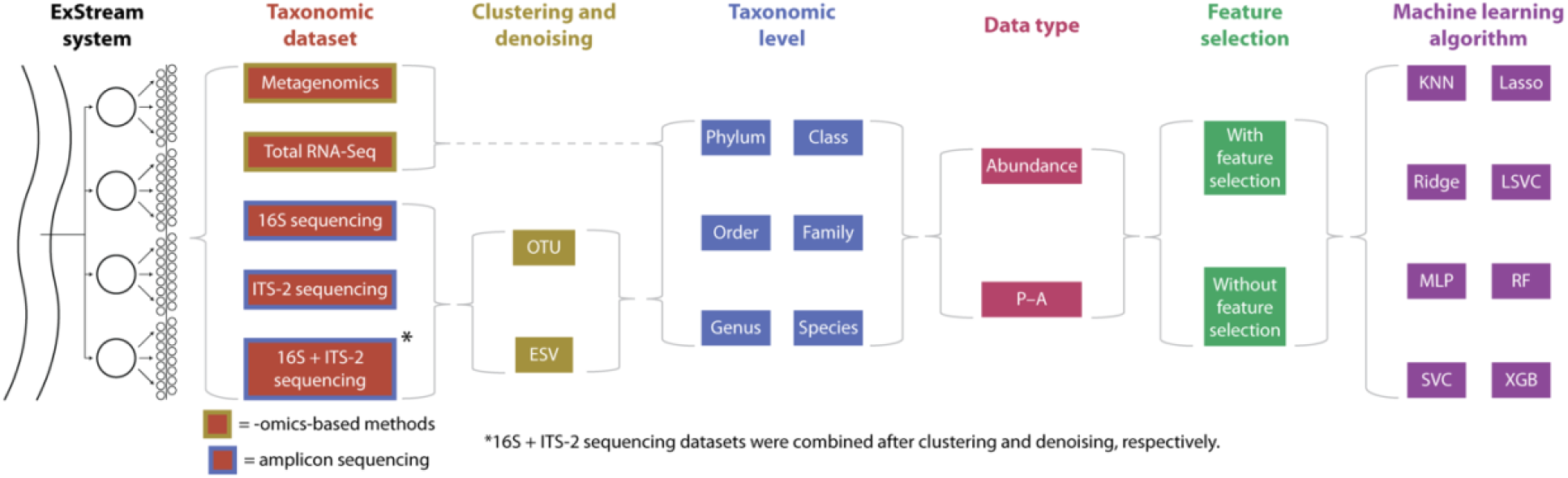
Summary of the study design. ExStream samples were processed with omics-based methods and amplicon sequencing, and HTS data were processed using two clustering methods (only applicable to amplicon sequencing), six taxonomic levels, two data types, eight machine learning algorithms, and with or without feature selection, for a total of 1,536 evaluated combinations of sequencing and data-processing methods. KNN: k-Nearest Neighbors; Lasso: Logistic Lasso Regression, Ridge: Logistic Ridge Regression, LSVC: Linear Support Vector Classification, MLP: Multilayer Perceptron, RF: Random Forest, SVC: Support Vector Classification, XGB: XGBoost.

### 2.1 Experimental setup

#### 2.1.1 ExStream system

A detailed explanation of the ExStream system can be found in Mack *et al*. [49]. In summary, stream mesocosms were connected to the adjacent stream Bieber, which provided them with a constant water flow. The stream Bieber is part of the Rhine-Main-Observatory (https://deims.org/9f9ba137-342d-4813-ae58-a60911c3abc1), a Long-Term Ecological Research site [50,51]. Each mesocosm was set up using substrate and organisms from the stream. A random subset of the mesocosms was exposed to either the insecticide chlorantraniliprole (Coragen, DuPont), increased fine sediment concentration, or both. Both insecticides and fine sediment are known key stressors of aquatic environments introduced into streams by agricultural runoff. The stressors were induced using a 4×2 factorial design by adding 0.2 μg/L, 2 μg/L, and 20 μg/L (acute stressor phase, 4 days) or 0.02 μg/L, 0.2 μg/L, and 2 μg/L (reduced stressor phase, 17 days) of the insecticide and 450 mL of fine sediment (< 2 mm) to the mesocosms. Each possible combination of stressor levels was replicated eight times in addition to eight control mesocosms that did not receive any stressor, resulting in 64 mesocosms.

#### 2.1.2 Assessment of microbial community compositions

The goal of the ExStream experiment was to evaluate the individual and combined effects of the applied stressors on biodiversity and organic matter decomposition in streams. To investigate organic matter decomposition, cotton strips were added to all mesocosms. Cotton strips are mainly made of cellulose, which is a major source of carbon in stream ecosystems. Therefore, analyzing the biofilm on the cotton strips allowed the analysis of the diversity of microbial communities degrading organic matter.

The experiment was divided into a colonization phase (days -21 to -1) and a stressor phase (days 0 to 21). Two cotton strips were added to each of the 64 mesocosms on day -17 (128 in total) and recovered after 28 or 35 days, respectively (for more information on the phases and cotton strip addition and recovery see Mack *et al*. [49]). Four cotton strips were washed away during the experiment, so 124 cotton strips were recovered in total. A 2-cm-long piece of each cotton strip was cut off and transferred into a ZR BashingBead Lysis Tube (0.1 & 0.5 mm) pre-filled with 1 mL of DNA/RNAShield (Zymo Research, Freiburg, Germany) using sterile laboratory gloves, forceps, and scissors. The samples were transferred to a laboratory, stored at -20 °C, and then homogenized using a bead mill homogenizer (MM 400, Retsch, Haan, Germany) at 1,800 rpm for 30 minutes. 300 μL of each lysate were processed for amplicon sequencing at the University of Duisburg-Essen, Germany, and the remainder of the lysates was shipped to the University of Guelph, Canada, on dry ice and processed for metagenomics and total RNA-Seq.

### 2.2 Laboratory processing

#### 2.2.1 Laboratory processing of amplicon sequencing

Amplicon sequencing was carried out following the workflow described by Buchner *et al*. [52]. All subsequent processing steps were completed on a Biomek FX^P^ liquid handling workstation (Beckman Coulter, Brea, CA, USA). Briefly, replication of the samples was carried out before DNA extraction by transferring 60 μL from the bead-beating tubes to deep-well plates pre-filled with 133 μL of TNES buffer (50 mM Tris, 400 mM NaCl, 100 mM EDTA, 0.5% SDS, pH 7.5) and 6 μL of Proteinase K (10 mg/mL) following incubation for 3 hours at 55 °C for complete lysis of the samples. DNA was extracted using a modified version of the NucleoMag Tissue kit (Macherey Nagel, Düren, Germany; for modifications see Buchner *et al*. [52]. Extraction success was verified using a 1% agarose gel.

The PCR for the amplicon library was performed using a two-step PCR protocol following Zizka *et al*. [53]. Samples were amplified in a first-step PCR using the Qiagen Multiplex Plus Kit (Qiagen, Hilden, Germany) with a final concentration of 1x Multiplex Mastermix, 200 mM of each primer (515F & 806R for 16S [54] and ITS3-CS1 & ITS4-CS2 for ITS-2 [55]), and 1 μL of DNA, and filled up to a total volume of 10 μL with PCR-grade water. The amplification protocol was: 5 min of initial denaturation, 25 cycles of 30 s denaturation at 95 °C, 90 s of annealing at 50 °C for 16S and 55 °C for ITS-2, and 30 s of extension at 72 °C, finished by a final elongation step of 10 min at 68 °C. For subsequent demultiplexing, each of the PCR plates was tagged with a unique combination of inline tags (Supplemental File S1).

The first-step PCR results were cleaned up with magnetic beads. The PCR product was mixed with clean-up buffer (2.5 M NaCl, 10 mM Tris, 1 mM EDTA, 20% PEG 8000, 0.05% Tween 20, 2% carboxylated Sera-Mag SpeedsBeads (Cytiva Life Sciences, Marlborough, MA, USA), pH 8) at a 0.8x ratio and incubated for 5 minutes, washed two times with wash buffer (10 mM Tris, 80% EtOH, pH 7.5) for 30 s, dried for 5 minutes at RT and finally eluted in 40 μL of elution buffer (10 mM Tris, pH 8.5).

During the second-step PCR, samples were amplified with a final concentration of 1x Multiplex Mastermix, 1x Coralload Loading Dye, 100 mM of each primer, and 2 μL of the first-step product. Cycling conditions were the same as in the first-step PCR except for 61 °C as annealing temperature and a decreased cycle number of 20. In the second-step PCR, each of the 96 wells was individually tagged so that the combination of the in-line tag from the first-step PCR and the index-read of the second-step PCR yielded a unique combination per sample. PCR success was verified using a 1% agarose gel.

PCR products were normalized to equal concentrations with normalization buffer (same as clean-up buffer, but with only 0.1% beads) following the same protocol as the clean-up after the first step but with a ratio of 0.7x and an elution volume of 50 μL. All normalized products were pooled in the final libraries in equal parts. The libraries were concentrated using a silica-membrane spin column (Epoch Life Science, Missouri City, TX, USA) by mixing 1 volume of the library with 2 volumes of binding buffer (3 M Guanidine Hydrochloride, 90 EtOH, 10 mM Bis-Tris, pH 6) for the binding step (1 min centrifugation, 11,000 x g), 2 washing steps (30 s centrifugation, 11,000 x g) with wash buffer and a final elution (3 min incubation at RT, followed by 1 min centrifugation at 11,000 x g) with 100 μL elution buffer. Library concentrations were quantified on a Fragment Analyzer (High Sensitivity NGS Fragment Analysis Kit; Advanced Analytical, Ankeny, USA). The libraries were then sequenced using the Illumina MiSeq platform with 2 lanes for each library with a paired-end kit (V2, 2×250 bp for 16S and V3, 2×300 bp for ITS) at CeGat (Tübingen, Germany).

#### 2.2.2 Laboratory processing of metagenomics and total RNA-Seq

DNA and total RNA were separately extracted from samples in 96-well plates using the NucleoMag DNA/RNA Water kit (D-MARK Biosciences, Toronto, Canada) that includes magnetic beads. Instead of using a magnetic plate to separate magnetic beads from buffers, we used the Magnetic Bead Extraction Replicator (V&P Scientific, San Diego, U.S.A.), which allows for the transfer of all magnetic beads from one lysate/buffer/elution plate to another without the need to remove the supernatant from individual wells (for the modified protocol, see <u>dx.doi.org/10.17504/protocols.io.bp2l69n2dlqe/v1</u>). The RNA extraction protocol involved a 25-min-long rDNase incubation step to digest DNA. Since the 96-well plates were open during the entire extraction, which posed a contamination risk, we added one negative extraction control to each row on each plate by replacing lysate with pure water. All extractions were performed under a sterile hood. DNA/RNA concentrations of all extracts and all negative extraction controls were measured using a Qubit fluorometer with the dsDNA HS Assay Kit and the RNA HS ASSAY Kit, respectively (Thermo Fisher Scientific, Burlington, Canada).

DNA and RNA libraries of all samples and negative extraction controls were prepared for metagenomics and total RNA-Seq using the NEBNext Ultra II DNA Library Prep Kit for Illumina and the NEBNext Ultra II Directional RNA Library Prep Kit for Illumina, respectively (New England Biolabs, Whitby, Canada). For RNA library preps, we did not perform mRNA enrichment or rRNA removal and instead processed the entire RNA. The RNA library prep kit has a default insert size of 200 bp, and we chose an insert size of 150–350 bp for the DNA library preps to keep insert sizes approximately consistent. After library prep, we randomly selected 8 DNA sample libraries, 3 negative DNA extraction control libraries, 7 RNA sample libraries, and 4 negative RNA extraction control libraries and sent 2.5 μL of each to the AAC Genomics Facility at the University of Guelph, Canada for analysis on an Agilent Bioanalyzer 2100 system (Agilent Technologies, U.S.A.) to confirm successful library preps and check for contaminations in negative extraction control libraries. After consultation with the sequencing facility (Centre for Applied Genomics, Hospital for Sick Children, Toronto, Canada), we cleaned up all DNA and RNA libraries following the DNA/RNA library prep kit manual to remove primer dimers and unincorporated primers.

We pooled 5 μL of each DNA and RNA library for sequencing, respectively, including negative extraction controls. We pooled equal volumes instead of equal concentrations because this pooling strategy allows for an equal relative sequencing depth per sample as opposed to an equal total sequencing depth. That way, the relative number of reads per sample mirrored the relative amount of DNA/RNA, avoiding an over- or underrepresentation of samples with higher or lower DNA/RNA amounts. Size distributions of the DNA and RNA library pools were assessed with a bioanalyzer by the sequencing facility, and the average fragment size was 386 bp for the DNA library pool and 436 bp for the RNA library pool. Both pools were paired-end (2×100 bp) sequenced in a 50:50 ratio on a single lane of a NovaSeq 6000 SP flowcell.

### 2.3 Bioinformatics

#### 2.3.1 Bioinformatics of amplicon sequencing

Raw data of the sequencing runs were delivered demultiplexed by index reads. Further demultiplexing by inline tags was done with the Python script “demultiplexer” (v1.1.0, https://github.com/DominikBuchner/demultiplexer). Sequences were subsequently processed with APSCALE v1.4 [56] using default parameters. Paired-end reads were merged using vsearch v2.21.1 [57]. Primer sequences were trimmed with cutadapt v3.5 [58]. For 16S sequencing, only sequences with a length of 252 ± 10 bp were retained, and for ITS-2 sequencing, only sequences with a length ranging from 240 to 460 bp were retained. Only sequences with an expected error of 1 passed quality filtering. Reads were dereplicated and singletons were removed. For OTU generation, sequence clustering was performed with a similarity threshold of 97%, and for ESV generation, denoising was carried out with an alpha value of 2 and a minimum size of 8 as implemented in vsearch. Before taxonomic assignment, the resulting OTU and ESV tables were filtered for potentially biased sequences with the LULU algorithm [59] implemented in APSCALE.

Subsequently, only reads found in both replicates of the same sample were summed up for all samples. After this initial data filtering, reads still left in the negative controls were subtracted from OTUs or ESVs, respectively, to generate final OTU and ESV tables. Taxonomic assignment was performed using DADA2 with default parameters in combination with the database SILVA 138.1 designed for DADA2 [60] for 16S sequences and the database UNITE [61] for ITS-2 sequences, respectively.

#### 2.3.2 Bioinformatics of metagenomics and total RNA-Seq

In an earlier study, we investigated 672 combinations of bioinformatic tools to identify the best-performing combination to process and taxonomically annotate microbial mock community datasets [31]. Based on these results, we processed both metagenomics and total RNA-Seq data as follows: we used Trimmomatic v0.39 [62] to trim the leading and trailing low-quality nucleotides of each read by cutting reads if the average quality of nucleotides in a sliding window of size 4 was below a PHRED score of 20. After trimming, we excluded reads shorter than 25 nucleotides and error-corrected reads using the error-correction module of the assembler SPAdes v3.14.1 [63]. Then we assembled the reads into scaffolds using MEGAHIT v1.2.9 [64] with the parameter ‘presets’ set to ‘meta-large’ to adjust k-mer sizes for the assembly of large and complex metagenomes. All other parameters were set to default. Subsequently, we mapped reads to assembled scaffolds to determine the abundance of each scaffold using BWA v0.7.17 [65] with default parameters. We processed mapped reads using the function *coverage* of samtools v1.10 [66] to obtain the mean per-base coverage for each scaffold. For taxonomic annotation, we used the SILVA132_NR99 SSU and LSU reference databases [67] in combination with kraken2 v2.1.1 [68] using default parameters. The setup of the kraken2 database for SILVA required manual adaptations, which are described in the Supplemental Material. All code utilized is available on GitHub (https://github.com/hempelc/metagenomics-vs-totalRNASeq).

### 2.4 Pre-processing of taxonomic data

The data were further processed in Python v3.7.9 [69]. The full code is available on GitHub (https://github.com/hempelc/exstream-metagenomics-totalrnaseq-ml) and uses the modules Pandas v1.3.5 [70] and NumPy v1.21.3 [71]. We trained and evaluated machine learning models based on phylum, class, order, family, genus, and species to assess differences in SPP among taxonomic levels. Because both metagenomics and total RNA-Seq datasets consisted of mean per-base coverage while amplicon sequencing datasets consisted of absolute read counts, we employed two different approaches to determine taxa abundances for each taxonomic level. When aggregating metagenomic and total RNA-Seq taxonomic datasets for each level separately, we selected all scaffolds assigned to each detected taxon and determined each taxon’s per-base coverage as a proxy of abundance as follows:

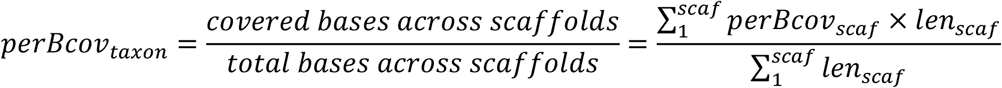

where *perBcov*_*taxon*_ represents the per-base coverage of a taxon, *scaf* represents the number of scaffolds assigned to a taxon, and *perbcov*_*scaf*_ and *len*_*scaf*_ represent the per-base coverage and length of each scaffold.

This procedure ensured that the per-base coverage of metagenomic and total RNA-Seq taxa reflected their cumulative scaffold length. We then converted absolute per-base coverages into relative per-base coverages, i.e., relative abundances.

When aggregating abundances based on amplicon sequencing data for each taxonomic level separately, we determined taxa abundances as the cumulative read count of each detected taxon and turned absolute abundances into relative abundances.

For metagenomics and total RNA-Seq samples, negative extraction controls were subtracted from samples that were co-extracted with the controls. Therefore, we converted relative abundances into absolute abundances by multiplying relative abundances by the number of reads per sample, summarized the absolute abundances of taxa among all negative extraction controls per plate, and subtracted the cumulative absolute abundance of each taxon detected within controls from the actual samples of the same plate. Afterwards, we turned absolute abundances back into relative abundances.

We then excluded the taxonomic entry *NA* from all datasets, which represented the relative abundance of sequences that could not be taxonomically annotated, likely due to missing references in databases or sequencing and data-processing errors. Next, we readjusted the relative abundances of all other taxa. In some datasets, some samples consisted only of sequences that could not be taxonomically annotated, meaning that they had a cumulative relative abundance of zero after excluding the *NA* entry. These samples were considered to have failed, and we excluded samples that failed in one dataset from all datasets to ensure that all datasets contained the same samples, which ultimately resulted in 121 samples per dataset.

To assess differences in SPP among data types, we evaluated abundance and P–A data. For P–A data, we set all relative abundances above 0 to 1 (0=absent, 1=present). For abundance data, we followed the appropriate steps for analyzing compositional data, as pointed out by Gloor *et al*. [72]. Therefore, we first applied simple multiplicative replacement to replace zeros among all relative abundances using the function *multiplicative_replacement* of the Python module scikit-bio v0.5.6 [73]. The function replaces zeros with a small positive value δ, which is based on the number of taxa while ensuring that the compositions still add up to 1. Then, we applied a centred log-ratio (clr) transformation using the function *clr* of scikit-bio, which captures the relationships between taxa and makes the data symmetric and linearly related. Since feature standardization is required by some machine learning algorithms, we further standardized taxa abundances using the function *StandardScaler* of the Python module scikit-learn v1.1.1 [74].

To include a multi-marker approach using both the ITS-2 and 16S marker genes in the evaluations, we combined the generated 16S and ITS-2 datasets by concatenating them using the clustering or denoising method (OTUs or ESVs). This resulted in eight taxonomic datasets that were evaluated (ITS-2 amplicon sequencing clustered into OTUs (ITS-2 OTU) or denoised into ESVs (ITS-2 ESV), 16S amplicon sequencing clustered into OTUs (16S OTU) or denoised into ESVs (16S ESV), multi-marker approach clustered into OTUs (16S + ITS-2 OTU) or denoised into ESVs (16S + ITS-2 ESV), metagenomics, and total RNA-Seq).

### 2.5 Biodiversity analysis

To analyze the biodiversity detected per taxonomic dataset, we determined the total number of detected taxa per taxonomic dataset, the number of unique taxa detected within only one taxonomic dataset, and the number of overlapping taxa between taxonomic datasets at the phylum and species level. Therefore, we translated all phyla and species names within 16S and ITS-2 datasets into NCBI taxonomy to match names across all datasets and utilized reference databases. We tested each name for matches with names in the scientific or non-scientific NCBI taxonomy (NCBI taxonomy file *names*.*dmp*, available through the NCBI archive as part of *taxdmp*.*zip*, https://ftp.ncbi.nlm.nih.gov/pub/taxonomy/), and if a match was found, the name was translated into the scientific NCBI name. If no match was found, we manually checked if the respective name was available on NCBI under a different scientific or non-scientific name, and if so, the alternative scientific name was used. Otherwise, the name was not available on NCBI and was used without translation. After translation, the number of overlapping phyla/species between taxonomic datasets was determined as the number of matches between the respective phyla/species within each taxonomic dataset, and the number of phyla/species unique to one taxonomic dataset was determined by subtracting the number of overlapping phyla/species from the total number of detected phyla/species.

### 2.6 Machine learning

#### 2.6.1 Data preprocessing

Taxon abundances/P–A represented independent features, and we defined the dependent feature as the combinations of applied insecticide level (none, low, medium, high) and fine sediment addition (normal fine sediment concentration, increased fine sediment concentration) for each sample, resulting in eight classes that were predicted by the machine learning algorithms. Since correlated independent features add noise, we removed correlated independent features by applying the SULOV (Searching for Uncorrelated List of Variables) algorithm using the function *FE_remove_variables_using_SULOV_method* of the Python module featurewiz v0.1.55 (https://github.com/AutoViML/featurewiz), which identifies all pairs of highly correlated independent features (features with a Pearson correlation coefficient of > 0.7 or < –0.7 by default), determines their Mutual Information Score (MIS) to the dependent feature, and keeps the independent feature with the highest MIS for each highly correlated feature pair.

#### 2.6.2 Test-train splitting and feature selection

Each ExStream mesocosm was sampled at two time points as part of the cotton strip assay, which meant that samples consisted of highly related sample pairs. When splitting the data sets into train and test sets, we ensured that highly related sample pairs always remained together in the train and test sets to avoid data leakage between the sets.

Initially, we applied a 90:10 train-test split to the datasets (109 train samples, 12 test samples) and performed training and testing without repetition, but due to large discrepancies between train and test scores, we changed the train-test split ratio to 80:20 (97 train samples, 24 test samples) and repeated both training and testing splits three times in total. During each repetition, we randomly selected 12 pairs (24 samples) of highly related samples for the test dataset and trained and tested all models across all datasets with the same randomly selected 12 sample pairs per repetition.

For feature selection, we used Recursive Feature Elimination to select the 20 most important features using the function *RFE* from scikit-learn with a *DecisionTreeClassifier* as the estimator.

#### 2.6.3 Model selection, training, and testing

It is generally recommended to test multiple machine learning algorithms [36], which is why we selected eight machine learning algorithms to predict stressor classes: k-Nearest Neighbors (KNN), Linear Support Vector Classification (LSVC), Logistic Ridge Regression (Ridge), Logistic Lasso Regression (Lasso), Multilayer Perceptron (MLP), Random Forest (RF), Support Vector Classification (SVC), and XGBoost (XGB). For thorough descriptions of these algorithms in a biological context see Greener *et al*. [36] and Ghannam & Techtmann [37].

All algorithms, except XGBoost, are available in scikit-learn. To run the XGBoost algorithm, we used the Python module xgboost v1.6.1 [75], which is compatible with scikit-learn. To optimize hyperparameters while avoiding overfitting, we performed Bayesian hyperparameter optimization with 10-fold cross-validation using the function *BayesSearchCV* of the Python module scikit-optimize v0.9.0 (https://github.com/scikit-optimize/scikit-optimize). The function is compatible with scikit-learn and builds a performance probability model for given hyperparameters, which is used to select the most promising hyperparameters through iterative performance evaluations. While not every possible hyperparameter combination is tested that way, this approach provides a good tradeoff between optimization results and runtime. Model prediction performance was evaluated using the Matthews Correlation Coefficient (MCC), which ranges from -1 to 1, where 1 means perfect predictions/performance, 0 means prediction performance as good as random guessing, and -1 means all predictions are wrong, and increments between -1 and 1 can be interpreted in the same way as increments of the Pearson correlation coefficient. All hyperparameters tested can be found in the publicly available code (https://github.com/hempelc/exstream-metagenomics-totalrnaseq-ml) and Supplemental File S2. The optimized hyperparameters were then used to train models on the entire training dataset, and model performances to predict classes of the testing dataset were evaluated using the MCC. During training on the entire dataset, learning curves were generated using the learning_curve function from scikit-learn. This process was repeated three times, as described above, and the mean average and standard deviation (SD) of the train and test MCC scores across the three repetitions were determined.

We tested each possible combination of taxonomic datasets (ITS-2, 16S, 16S + ITS-2, metagenomics, and total RNA-Seq), clustering or denoising methods (OTU, ESV; only applicable to amplicon sequencing data), taxonomic levels (phylum, class, order, family, genus, and species), data types (abundance, P–A), feature selection (with feature selection, without feature selection), and machine learning algorithms (KNN, Lasso, LSVC, Ridge, MLP, RF, SVC, and XGB), resulting in a total of 1,536 evaluated combinations.

### 2.7 Statistical analysis

We quantified the impact of sequencing types, taxonomic levels, data types, feature selection, and machine learning algorithms on SPP. For that, we converted all sequencing and data-processing methods into binary dummy variables and tested for significant correlations (*P* ≤ 0.05) between each sequencing and data-processing method and the test MCC by calculating Spearman’s rank correlation coefficient using the spearmanr function of the Python module SciPy v1.7.1 [76]. Additionally, we performed the same test for each sequencing type separately.

## 3 Results

### 3.1 HTS results

We obtained 248,707,817 paired-end reads from metagenomics (mean average per sample (M): 2M reads, standard deviation (SD): 2.4M reads), 206,096,238 from total RNA-Seq (mean average per sample: 1.7M reads, SD: 2.6M reads), 21,719,985 reads from 16S sequencing (mean average per sample: 152k reads, SD: 27k reads), and 27,033,469 reads from ITS-2 sequencing (mean average per sample: 214k reads, SD: 41k reads) (Figure S1; Bioproject number: PRJNA903104, SRA accession numbers: SRR22331748–SRR22332597). The SD of the mean average number of metagenomics and total RNA-Seq reads per sample was very high since we normalized metagenomics and total RNA-Seq libraries based on volume during library preparation so that the relative number of reads per sample mirrored the relative amount of DNA/RNA. This avoided an over- or underrepresentation of samples with higher or lower amounts of DNA/RNA but also led to substantial variations in the number of reads per metagenomics/total RNA-Seq library (Figure S1).

### 3.2 Biodiversity analysis

There were no taxa overlaps between ITS-2 and 16S sequencing at both the phylum and species level (Figure 2, for exact numbers, see Supplemental File S3), while either method had overlapping taxa with both metagenomics and total RNA-Seq. Metagenomics and total RNA-Seq shared more taxa with each other than with ITS-2 or 16S sequencing, especially at the species level. Metagenomics detected the most phyla (147) and species (3,713), but the number of species detected using metagenomics was much higher relative to that of other taxonomic datasets than the number of detected phyla. For total RNA-Seq, the number of detected phyla (87) was more than twice as high as that of ITS-2 (OTU: 23, ESV: 20) and 16S sequencing (OTU: 36, ESV: 32), and the number of detected species (927) was almost twice as high as that of ITS-2 sequencing (OTU: 677, ESV: 509) and much higher than that of 16S sequencing (OTU: 55, ESV: 116). 16S sequencing detected almost the same number of phyla as ITS-2 sequencing but by far the lowest number of species among all taxonomic datasets. In terms of taxa unique to one taxonomic dataset, metagenomics detected by far the most unique taxa. It detected more unique phyla (57) than the total number of phyla in ITS-2 or 16S sequencing and almost as many unique phyla as the total number of phyla in total RNA-Seq. Furthermore, metagenomics detected more unique species (2,796) than the total number of species detected by all other taxonomic datasets combined. The number of unique species in relation to the number of total species was low in total RNA-Seq in comparison to that of other taxonomic datasets. No unique phyla were detected with ITS-2 and 16S sequencing when either was denoised into ESVs. Within ITS-2 and 16S sequencing, OTU clustering and ESV denoising resulted in approximately the same number of taxa, but at the species level, the majority of detected taxa was unique, indicating a low species overlap between OTU clustering and ESV denoising.

**Figure 2:**
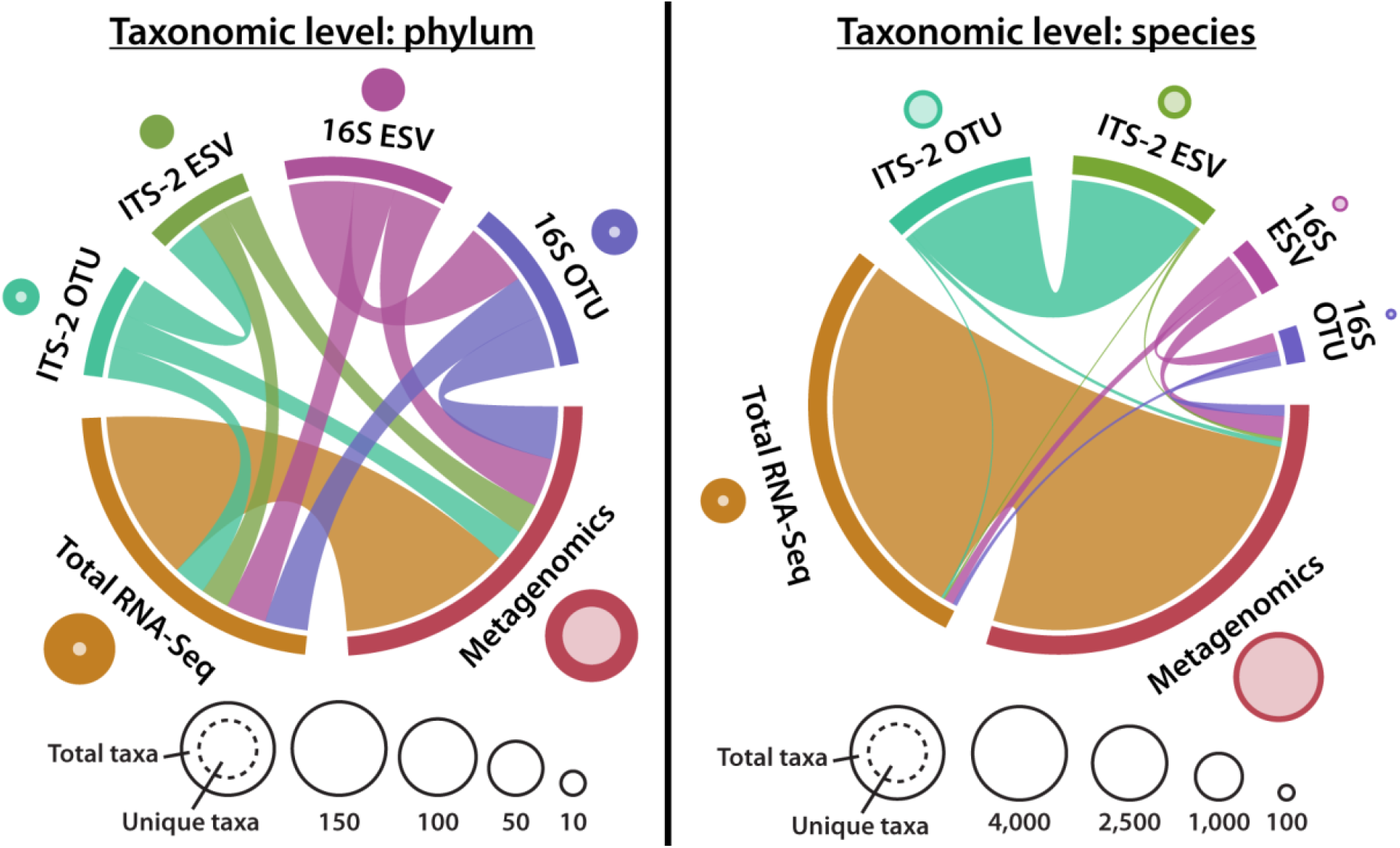
Number of total detected, unique, and overlapping taxa for each taxonomic dataset on the phylum and species level. The outer circle size next to each taxonomic dataset indicates the number of total detected taxa for that taxonomic dataset, and the inner circle size (faded color) indicates the number of unique taxa detected only for that taxonomic dataset. The chord diagram indicates the number of overlapping taxa between taxonomic datasets, where each outer bar represents the number of total overlapping taxa for each taxonomic dataset (also indicated by the non-faded circle area) and the width of connections between taxonomic datasets represents the number of overlapping taxa between taxonomic datasets. Metagenomics detected the most taxa overall (indicated by the large outer circle size) and the most unique taxa (indicated by the large inner circle size) for both taxonomic levels. Metagenomics and total RNA-Seq had a large overlap of taxa (indicated by the large width of connections between both taxonomic datasets), while 16S and ITS-2 sequencing had no overlaps with each other and smaller overlaps with both omics-based methods.

### 3.3 Impact of taxonomic datasets and data-processing methods on SPP

The MCC, i.e., SPP varied substantially across tested combinations of taxonomic datasets, clustering or denoising methods, taxonomic levels, machine learning algorithms, and feature selection (Figure 3; since data types had no significant impact on SPP (see Figures 4 and 5), only P–A-based SPPs are shown). MCC values ranged from below 0 (prediction SPP worse than random guessing) to 0.45 (moderate to good SPP). Feature selection overall improved SPP. ITS-2 sequencing and omics-based methods performed poorly overall, except for some combinations of ITS-2 sequencing with OTU clustering, whereas 16S sequencing and the multi-marker approach of combined 16S and ITS-2 markers performed better overall. The highest MCC of 0.448 was found for the following combination: 16S + ITS-2 sequencing, ESV denoising, genus level, P–A data, Lasso algorithm, with feature selection. For this combination, the learning curves generated during each training repetition indicated that the model was under-fitted, meaning that more data, i.e., more samples would have likely further increased SPP (Figure S2).

**Figure 3:**
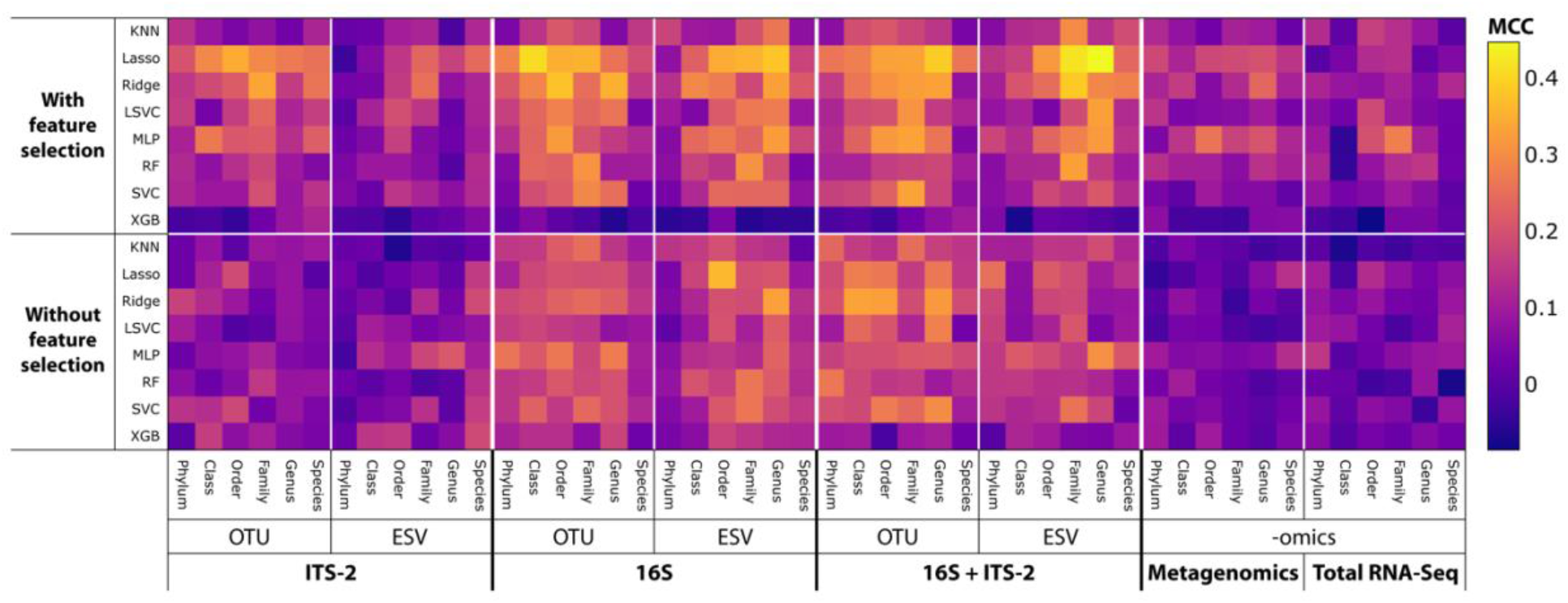
MCC, i.e., SPP across all combinations of sequencing and data-processing methods tested. Since data types had no significant impact on SPP (see Figures 4 and 5), only P–A-based SPPs are shown. SPPs varied substantially between combinations, indicated by the wide range and uneven distribution of MCC values.

**Figure 4:**
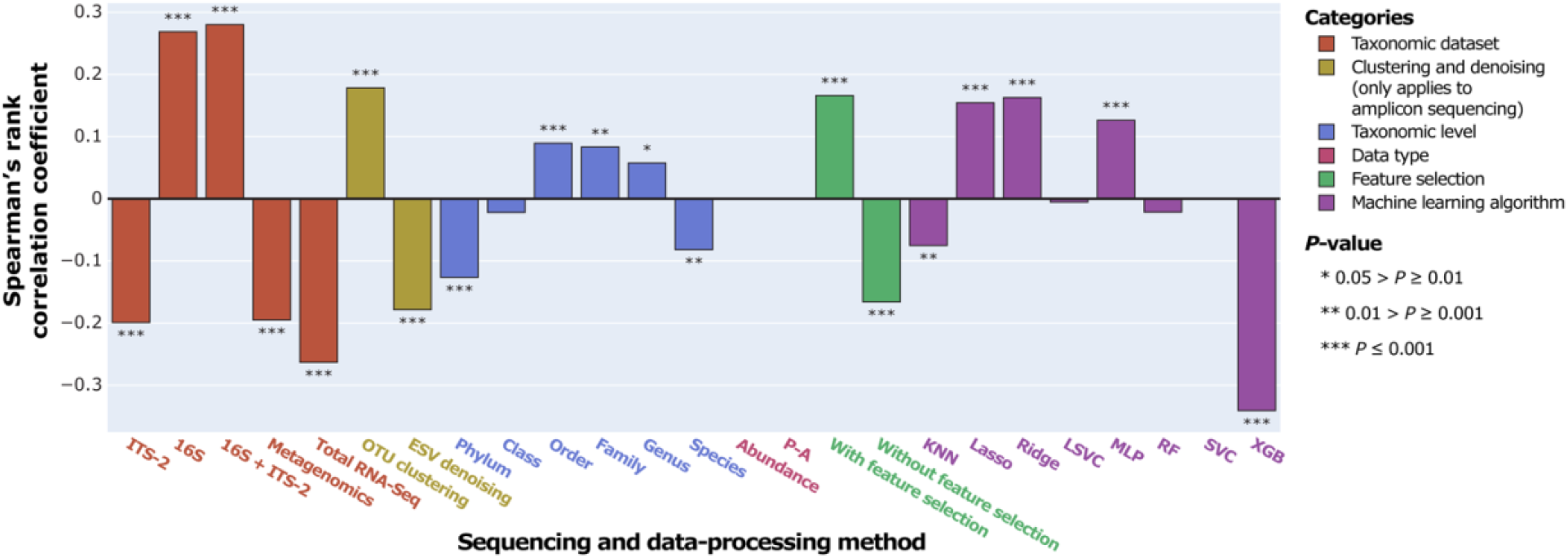
Correlation between MCC, i.e., SPP and sequencing and data-processing methods.

**Figure 5:**
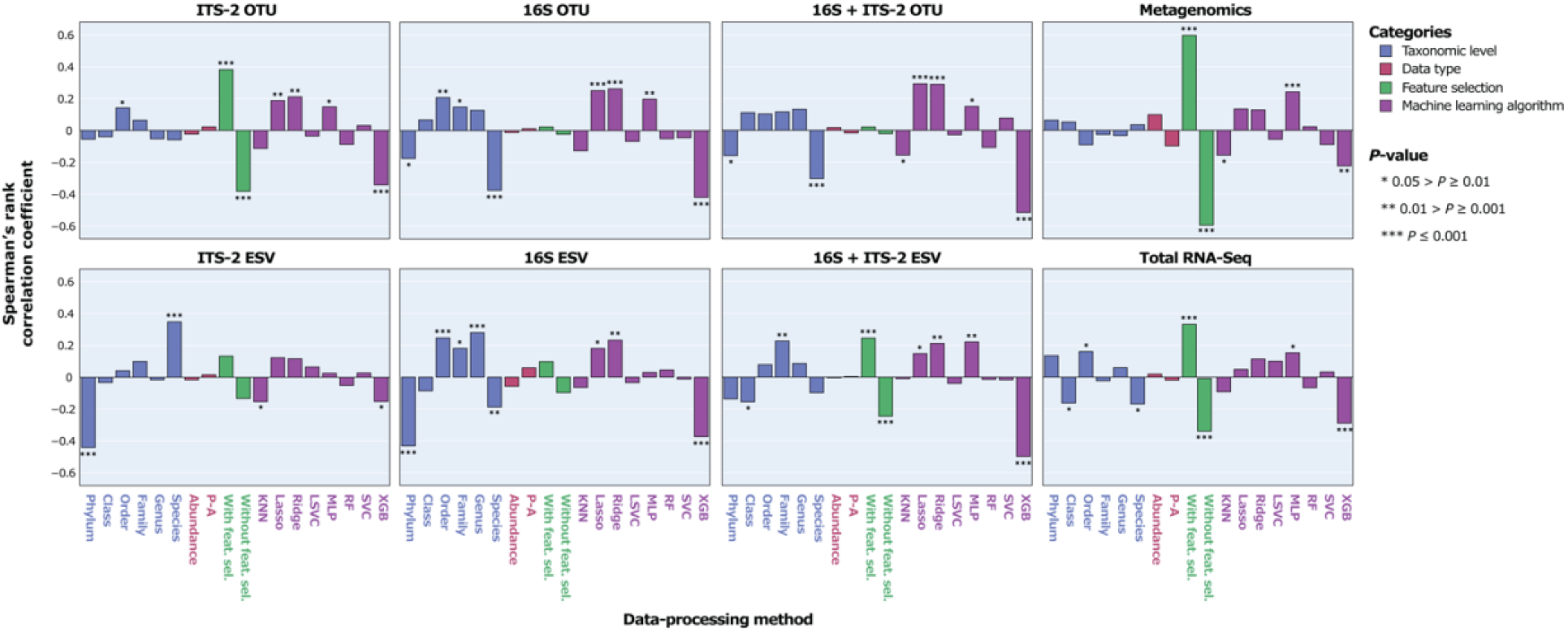
Correlation between MCC, i.e., SPP and data-processing methods for individual taxonomic datasets.

Overall, ITS-2 sequencing, metagenomics, and total RNA-Seq significantly negatively correlated with SPP, and 16S sequencing and combined 16S + ITS-2 markers significantly positively correlated with SPP (Figure 4). For amplicon sequencing, OTU clustering significantly increased SPP while ESV denoising significantly decreased SPP. Performance increased with increasing taxonomic resolution up to the order level and decreased at higher levels. Data types did not significantly correlate with SPP. Feature selection significantly increased SPP. SPPs varied between machine learning algorithms, with XGB performing by far the worst and Lasso and Ridge, which are both based on logistic regression, performing the best, followed by MLP.

The impact of data-processing methods on SPP varied between individual taxonomic datasets (Figure 5). For ITS-2 ESV, the species level was significantly positively correlated with SPP, which contrasted with all other taxonomic datasets. For metagenomics, no taxonomic level significantly correlated with SPP. Data types did not significantly correlate with SPP in any taxonomic dataset. Feature selection had the strongest impact on metagenomics and no impact on 16S OTU/ESV and 16S + ITS-2 OTU. Across all taxonomic datasets, XGB performed poorly. Lasso and Ridge performed significantly well for all taxonomic datasets except metagenomics, total RNA-Seq, and ITS-2 ESV (Ridge) and total RNA-Seq (Lasso). Overall, the impact of the data-processing methods was similar between 16S OTU/ESV, 16S + ITS-2 OTU/ESV, and ITS-2 OTU and differed between metagenomics, total RNA-Seq, and ITS-2 ESV.

## 4 Discussion

### 4.1 Biodiversity analysis

The number of total detected, unique, and overlapping taxa varied substantially between taxonomic datasets. ITS-2 and 16S sequencing had no taxa overlap, confirming that both markers were group-specific for bacteria and fungi, respectively. Metagenomics and total RNA-Seq had overlapping taxa with ITS-2 and 16S sequencing but also detected a high number of taxa that ITS-2 and 16S sequencing did not detect, confirming that omics-based methods cover groups across the tree of life, which is considered a major advantage over amplicon sequencing [21,24,77]. Many taxa found with total RNA-Seq were also found with metagenomics, but the latter also found an extremely high number of unique taxa. However, at the species level, ITS-2 and 16S sequencing detected a high number of unique taxa as well, showing that within target groups, amplicon sequencing had a higher resolution than omics-based methods. There is no clear consensus in the literature as to which method detects more taxa, with some studies showing that amplicon sequencing detects more taxa than omics-based methods [9,20], while others show that both methods detect equal amounts of taxa [77,78] or that omics-based methods outperform amplicon sequencing in terms of biodiversity coverage [21,24,79,80]. Biodiversity coverage also depends on how well an environment is represented in reference databases, and for less-studied environments that are poorly represented in reference databases, it is possible that the majority of omics-based sequences cannot be taxonomically annotated, resulting in low overall taxonomic resolution [9]. Our results support both hypotheses: (1) omics-based methods detect more taxa overall, and (2) amplicon sequencing detects more taxa within target groups, which aligns with the advantages and disadvantages of either approach. In theory, all taxa detected with amplicon sequencing should also have been detected with omics-based methods, but our results indicate that sequencing depth for omics-based methods must be increased substantially to be able to detect the same taxa. However, given continuous technological advancements in HTS capacities, sufficient sequencing depths should become more affordable, and in combination with the steady growth of reference databases, we expect omics-based methods to unilaterally detect more taxa than amplicon sequencing at equal or higher taxonomic resolution in the future.

### 4.2 Impact of sequencing methods on SPP

SPP varied substantially among taxonomic datasets. 16S sequencing was the only method positively correlated with SPP, and combining 16S with ITS-2 sequencing data slightly improved SPP. We expected omics-based methods to outperform amplicon sequencing, since they are not group-specific and can cover the entire tree of life, providing a more holistic picture of microbial communities; however, the opposite was the case, indicating that while omics-based methods did detect more taxa, they also generated more noise and/or data without correlation to stressors, which decreased SPP. This was further supported by the fact that the SPP of metagenomics, which detected the highest number of taxa, improved by far the most with feature selection, i.e., the exclusion of all but the 20 most relevant taxa for model performance. However, even with feature selection, metagenomics still showed poor overall SPP, indicating that the feature-selected taxa did not correlate with stressors or were poorly represented, likely because they were not sufficiently covered due to insufficient sequencing depth. Typical metagenomics experiments aim to generate between 1 and 10 Gb of metagenomic data per sample [81] while we generated on average 0.2 Gb metagenomic data per sample, which is one to two magnitudes lower. Based on our previous results showing that total RNA-Seq outperforms metagenomics in terms of identifying a microbial community and reconstructing SSU rRNA sequences [31,82], likely due to higher SSU rRNA sequence yield for total RNA-Seq, we expected that total RNA-Seq would perform better than metagenomics at low sequencing depth (on average 0.17 Gb total RNA-Seq data per sample). But total RNA-Seq performed even worse, so the sequencing depth of total RNA-Seq was likely insufficient as well.

Increasing the sequencing depth of omics-based methods to ensure that taxa with high bioindication potential are sufficiently represented might increase SPP but is currently also related to substantially higher costs. Almost all studies that utilize machine learning for taxonomically assigned HTS data in an environmental context involve amplicon sequencing [5,40,42–46], and to our knowledge, there is only one study that involves metagenomics in that context [47] and none that compare amplicon sequencing with omics-based methods. However, in a medical context, Marcos-Zambrano *et al*. [83] provide a thorough overview of human microbiome studies that utilize machine learning for HTS data, but while they list seven studies that applied machine learning to both amplicon sequencing and metagenomics data, only one of them compared the performance of both sequencing methods based on community composition [84], showing that amplicon sequencing outperformed metagenomics in classifying patients and the state of Crohn’s disease while metagenomics outperformed amplicon sequencing in classifying treatment response. These results further demonstrate that SPP is dependent on the environmental variables investigated. Multiple other medical studies utilizing machine learning for disease predictions based on metagenomics community compositions show good SPP for predicting colorectal cancer, inflammatory bowel disease, diabetes, rheumatoid arthritis, and liver cirrhosis [85–87]. These studies clearly show the potential of omics-based methods for medical applications, and further omics-based ecological research with sufficient sequencing depth is required to show if the methods hold the same potential for environmental stressor predictions.

### 4.3 Impact of data-processing methods on SPP

Data-processing methods had a substantial impact on SPP, and based on the utilized methods, SPP could range from low to high within one taxonomic dataset.

#### 4.3.1 Impact of clustering and denoising methods on SPP

For amplicon sequencing data, OTU clustering significantly improved SPP while ESV denoising significantly decreased SPP. This observation is in contrast to the emerging recommendation to denoise amplicon sequences into ESVs [88,89]. Studies comparing OTU clustering and ESV denoising approaches did not yet reach a consensus, showing that either both approaches lead to similar results [90–92], ESV denoising outperforms OTU clustering [93–95], or vice versa [96,97]. Our results support the latter, although more similar studies are required to determine if clustering or denoising is more appropriate for machine-learning-based environmental predictions based on microbial communities.

#### 4.3.2 Impact of taxonomic levels on SPP

In theory, a higher taxonomic resolution should provide a better picture of microbial communities, but our results show that the species level correlated worse with SPP than the genus, family, order, and even class levels. For ITS-2 sequencing and omics-based methods, the high number of detected taxa at the species level might have added more noise than value to the data, which reduced SPP. This is indicated by the significantly positive impact of feature selection on SPP, i.e., the limitation of the number of included taxa. However, for 16S sequencing, feature selection had no impact on SPP while the species level still negatively correlated with SPP, which requires an additional explanation. This result may be related to the number of sequences that could not be assigned to the species level and were consequently dropped. The higher the taxonomic level considered, the harder it is to annotate taxonomy due to the lack of reference sequences in databases and the more sequences are dropped from the downstream analysis. In microbiome amplicon sequencing studies, the taxonomic resolution is usually limited to the genus level due to the difficulty in designing primers that resolve microbial communities at the species level [88]. Metagenomics allows for taxonomic resolutions at the species level or even strain level, but this requires sufficient sequencing depth [88]. Dropping sequences from the analysis is equivalent to a loss of information, which could decrease SPP. It is also possible that correlations between taxa and environmental variables are higher at lower taxonomic levels because lower taxonomic groups can be overall ecologically coherent, i.e., share similar physiologies, while higher taxonomic groups can be ecologically incoherent and have very different physiologies [98–100]. Once reference databases have been extensively expanded and most sequences can be taxonomically annotated, it will be possible to determine if the lack of reference sequences or ecological incoherency of species explains lower SPP at the species level.

#### 4.3.3 Impact of data types on SPP

We were surprised that the data types (abundance/P–A) did not have an impact on SPP, given that many studies focus on methods to improve abundance estimates from HTS data [72,101–103]. The difference in abundance and P–A data lies in the weight of the taxa; in P–A data, abundant and rare taxa are weighted equally, making the data more sensitive to noise but also to subtle differences in community composition. Using simulated data, Koh *et al*. [104] demonstrated that P–A data is more powerful when taxa associated with an environmental variable are rare while abundance data is more powerful when those taxa are abundant. However, a large-scale morphological study on benthic invertebrates showed that ecological status classifications based on abundance and P–A data showed only minor variations [105]. In a microbial context, multiple HTS studies showed similar correlations of both abundance and P–A data with environmental variables [106–108], while some studies showed that correlations differed between data types [109,110]. These results indicate that the impact of data types might depend on the studied environmental variables, but if further research shows that both data types have similar predictive power for environmental assessments, as our results suggest, then P–A data could be used exclusively in future environmental assessment studies. That way, the rather complex and partially disagreeing statistical methods required when working with compositional data, i.e., HTS abundance data could be avoided [72,101–103]. Furthermore, if abundance and P–A data generate similar results, then the often-cited advantage of metagenomics to generate abundance data without bias resulting from target PCR [88,111] would become irrelevant, which would decrease the value of omics-based approaches in comparison to amplicon sequencing.

#### 4.3.4 Impact of feature selection on SPP

Feature selection can be applied to microbial data to remove noninformative, noisy, or redundant features [37]. This is generally recommended because the high number of observed features can increase the risk of overfitting, which is described as the “curse of dimensionality” [112]. However, feature selection goes against the proposed idea that a more holistic picture of environmental microbial communities is beneficial for predicting environmental variables as it reduces the number of taxa included in prediction models. Our results suggest that feature selection improves SPP overall and especially for metagenomics, while the SPP of 16S sequencing was not impacted by feature selection. This indicates that the increased biodiversity coverage of omics-based methods might in fact not be beneficial for machine learning predictions and that datasets covering a lower number of taxa, as generated by amplicon sequencing, might result in more accurate and precise predictions. It should be noted, though, that ITS-2 sequencing detected approximately as many species as total RNA-Seq, and feature selection did increase the SPP of ITS-2 sequencing, showing that amplicon sequencing can be impacted by feature selection. Furthermore, the sequencing depth of metagenomics and total RNA-Seq in our study was very low, which could have influenced the impact of feature selection. If similar studies with a sufficient sequencing depth come to the same conclusion that omics-based methods in fact detect too many taxa for accurate and precise environmental assessments and require feature selection, then this would strongly tip the balance in favor of amplicon sequencing.

#### 4.3.5 Impact of machine learning algorithms on SPP

Machine learning algorithms had a substantial impact on SPP, and even when applying two different algorithms to the same data set, the resulting MCC could range from 0.38 to -0.05. This illustrates the importance of testing multiple machine learning algorithms, which is recommended in general [36]. One of the most commonly applied machine learning classification algorithms for HTS data is RF [5,37,43,44,46,83,113], which reveals which feature contributed most to a prediction. Other popular algorithms are XGB, Support Vector Machines (which include SVC and LSVC), Logistic Regression, and KNN [36,37,83]. However, among those algorithms, RF and (L)SVC did not significantly correlate with SPP in our study, while XGB and KNN significantly negatively correlated with SPP and only logistic regression, specifically Lasso and Ridge, significantly positively correlated with SPP. Linear algorithms have the lowest flexibility among all popular machine learning algorithms, and while the higher flexibility of other algorithms is considered beneficial for the analysis of large and complex data, this was not the case for our study. In contrast, MLP, which represents a simple neural network (NN) with the highest flexibility among all algorithms tested in our study, performed overall the best after Lasso and Ridge and specifically the best for omics-based methods that generated the largest datasets. NNs are currently among the most powerful machine learning algorithms for the analysis of extremely large data, and their impact is so significant that an entirely new field of research emerged around NNs, called deep learning [36]. To unfold their potential, NNs require large amounts of samples that usually go beyond the number of samples generated in a single biological study. However, thousands of sampling sites are monitored for routine environmental assessments, and once sufficient sequencing depths of omics samples become more affordable, it will be interesting to see if NNs are required for good SPP based on omics data or if less complex machine learning algorithms will be sufficient or even more appropriate.

Overall, our study shows that data-processing methods should be chosen carefully since they can have a high impact on SPP and that methods resulting in the single best SPP are not necessarily the most appropriate overall. Therefore, we conclude that it is advisable to explore multiple sequencing and, in particular, data-processing methods to maximize prediction performance.

### 4.4 Perspectives for ecological assessments

The highest MCC, i.e., the best SPP observed in our study was 0.448, indicating moderate to good performance. While this is promising, stressor predictions must be more accurate and precise to reach the standard for applied ecological assessments. However, while the stressors tested in our study (insecticide and increased fine sediment deposition) have direct negative effects on typical indicator organisms (e.g., benthic macroinvertebrates), little is known about their effects on microbial communities. Since many microbes are a good indicator of ecosystem health and respond sensitively to stressors, we expected a shift of the microbial communities under exposure to insecticide and increased fine sediment deposition, at least due to indirect top-down effects caused by the reduced abundance of benthic macroinvertebrates that typically graze on cotton strips. But it is also possible that direct or indirect effects of the stressors on microbes were too low to cause a sufficient shift in microbial communities for taxonomy-based stressor predictions or even that increased fine sediment deposition was beneficial for microbial communities because it provided additional surface habitat for microbes or stimulated organic matter decomposition through physical abrasion of the cotton strips. Therefore, our observed insufficient SPP could also be a consequence of stressor choice rather than limitations of machine learning, especially since other studies show good performance of machine learning models for environmental assessments based on amplicon sequencing [40,42–45]. Smith *et al*. [5] showed that the performance of prediction models can highly vary based on the predicted environmental variables (including stressor variables). When they attempted to predict 38 geochemical groundwater variables based on 16S sequencing data, the predicted and actual values of 26 variables significantly correlated with each other while those of 12 variables did not. This was further supported by Hermans *et al*. [46], who predicted seven soil variables based on 16S sequencing data, and the correlations between predicted and actual values ranged from weak to strong and were further dependent on the land use type of the investigated samples. This raises the need for more exploratory research using different stressors until machine learning can be broadly applied to ecological assessments that involve many stressors. Nevertheless, the learning curves generated for our best model indicate that more samples likely would have increased SPP. This result is promising because it shows that further sampling likely would have revealed subtle yet distinctive community shifts that would have allowed for better predictions without requiring further knowledge about the direct or indirect effects of the stressors on microbes, which further highlights the potential of machine learning for HTS-based environmental assessments given sufficient sampling size.

We have only investigated the taxonomic information generated by metagenomics and total RNA-Seq, but both methods also generate information on functional diversity (metagenomics) and functional gene expression (total RNA-Seq). This information can also be integrated, which is why omics-based methods are gaining increased attention for environmental assessments [7,32–34], and it remains to be seen to what extent SPP can be increased by integrating taxonomical and functional information.

## 5 Conclusions

We demonstrate that both sequencing and data-processing methods have a substantial impact on environmental stressor prediction when applying machine learning to taxonomically assigned HTS data. Omics-based methods detected much more taxa than amplicon sequencing, and while this is considered an advantage, amplicon sequencing, specifically 16S sequencing, outperformed all other sequencing methods in terms of stressor prediction performance (SPP). However, the best observed SPP for 16S sequencing was only moderate to good, meaning that further improvements are necessary to meet the required standard for applied ecological assessments. Nevertheless, learning curves indicated that more samples would likely have increased SPP, demonstrating the potential for further research. Omics-based methods performed poorly, but this was likely due to insufficient sequencing depth, and given that other studies demonstrated the potential of omics-based methods in combination with machine learning, further omics-based ecological research with sufficient sequencing depth is required to show if this approach holds the same potential for environmental stressor predictions. Data types had no impact on SPP while feature selection significantly improved SPP for omics-based methods but not for amplicon sequencing, and if similar studies confirm these results, then this would strongly favor the application of amplicon sequencing over omics-based methods for environmental assessments. However, we only investigated taxonomic information, but omics-based methods also generate functional information, and it remains to be tested whether the integration of taxonomic and functional information can further improve omics-based environmental assessments.

## 6 Declarations

### 6.1 Availability of data and materials

The sequencing data has been deposited in the NCBI Sequence Read Archive (SRA) and is available under Bioproject number PRJNA903104 and SRA accession numbers SRR22331748– SRR22332597.

### 6.2 Competing interests

The authors declare that they have no competing interests.

### 6.3 Funding

CAH was funded through the Canada First Research Excellence Fund to the program CFREF - Food from Thought at the University of Guelph. DB, MVB, and the field experiment were funded through the DFG grants LE 2323/9-1, MA, and SCHA. LM was funded through the Land2Sea project (Aquatic Ecosystem Services in a Changing World, https://land2sea.ucd.ie/; funded under the Joint BiodivERsA-Belmont Forum call and the DFG) and the DFG project LE2323/9-1 / MA XXXXX 418091530.

### 6.4 Authors’ contributions

DB, LM, MVB, and FL designed the experiment. DB, LM, and MVB conducted the experiment and collected the samples. DB, LM, and CAH processed the samples. DB and CAH processed the sequencing data. CAH and DT analyzed the data. CH drafted the manuscript. All authors read and approved the final manuscript.

## 6.5 Acknowledgements

We are grateful to Christoph Mayer, Peter Haase, and Ralf Schäfer for their support during the grant application, and we thank Verena Schreiner for performing the pesticide analysis and Romana Salis for performing the taxonomic annotation of amplicon sequencing data.

## 8 Supplemental Figures

**Supplementary Figure 1:**
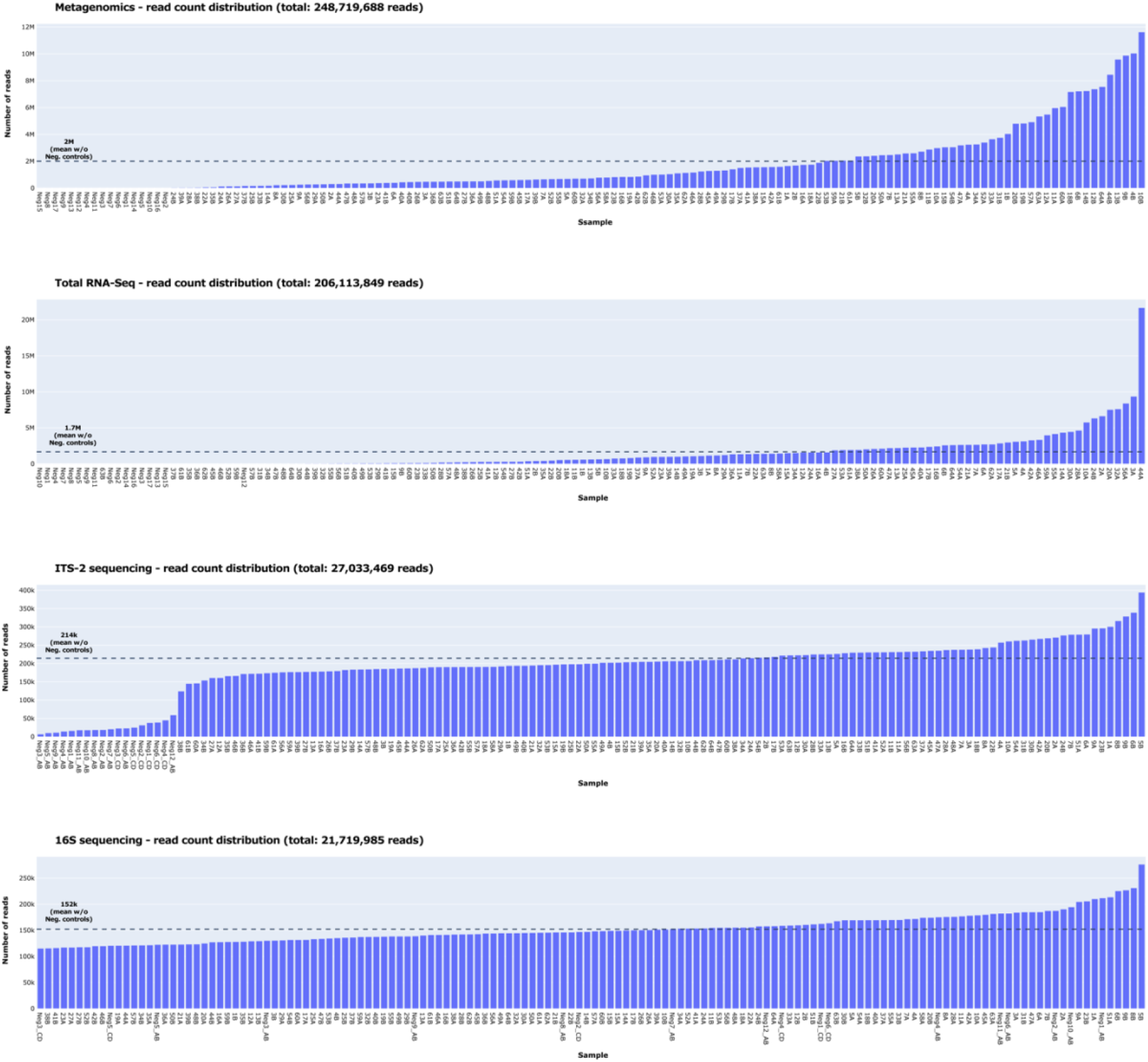
Number of reads and read distribution obtained for metagenomics, total RNA-Seq, 16S sequencing, and ITS-2 sequencing. Mean average numbers of reads (excluding negative controls) are indicated by dashed lines. Note that the y-axis range differs among graphs. The number of reads received per metagenomics and total RNA-Seq sample varied substantially since libraries were normalized based on volume rather than concentration so that the relative number of reads per sample mirrored the relative amount of DNA/RNA, avoiding an over- or underrepresentation of samples with higher or lower amounts of DNA/RNA.

**Supplementary Figure 2:**
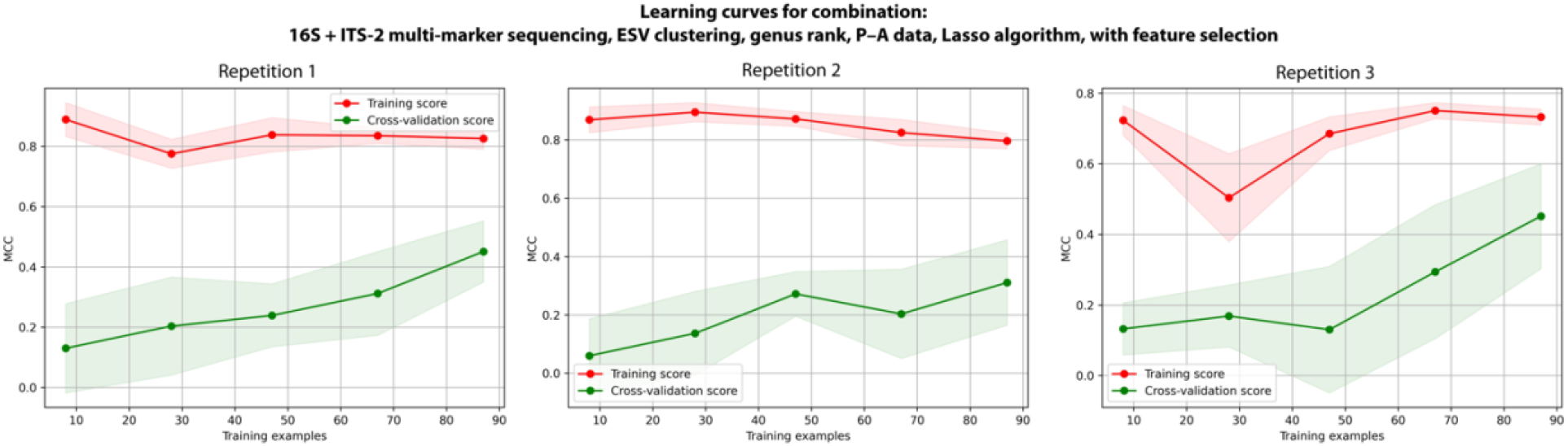
Learning curves generated during model training of the best-performing combination of sequencing and data-processing methods across three repetitions. The constant increase in the cross-validation score indicates that the models were underfitted.

## 9 Supplemental material

### 9.1 Setup of the SSU and LSU SILVA kraken2 database

We downloaded the available SILVA132_NR99 SSU and LSU databases and merged them into one SILVA database, removing duplicates that falsely occurred in both databases according to the SILVA-arb support. We standardized the taxonomy nomenclature of the SILVA database by translating the taxonomy of all SILVA reference sequences into the NCBI Genbank taxonomy before building SILVA reference databases. Note that some SILVA reference sequences are annotated with species names that do not match the rest of their taxonomic annotation. To translate the SILVA taxonomy into Genbank taxonomy, we first made sure that the last taxonomic level of each SILVA reference sequence was a species name that fit the remaining taxonomy. Therefore, we checked if the genus part of the species name matched the genus name for each SILVA reference sequence, and if not, the species name was converted to *NA*. Then, we checked each taxonomic level of each SILVA reference sequence for matches among first scientific and then non-scientific names in the Genbank taxonomy file names.dmp (available through the NCBI archive, part of taxdmp.zip, https://ftp.ncbi.nlm.nih.gov/pub/taxonomy/), beginning at the last level, and if a match was found, the respective Genbank taxonomic ID was used for that SILVA reference sequence. If no match was found for that level, the process was repeated for the next higher level until a match was found. If no match was found overall, the reference sequence was assigned with *NA*. Note that we removed all Genbank names containing “environmental”, “uncultured”, “unidentified”, and “metagenome” from the names.dmp file prior to translation to exclude them from the matching process. The kraken2 database was set up following the developer’s instructions (https://github.com/DerrickWood/kraken2/blob/master/docs/MANUAL.markdown). To set up the kraken2 database for the merged SSU and LSU SILVA database, we translated the taxonomy of each reference sequence into Genbank taxonomy IDs as described above, added the IDs to each reference sequence, and created files in the same format as the taxonomy files of the Genbank database, names.dmp and nodes.dmp (part of taxdmp.zip, https://ftp.ncbi.nlm.nih.gov/pub/taxonomy/). We automated the entire process with a script that is available on GitHub (https://github.com/hempelc/metagenomics-vs-totalRNASeq), which incorporates the SILVA taxonomy files taxmap_slv_ssu_ref_nr_138.1.txt.gz, taxmap_slv_lsu_ref_nr_138.1.txt.gz, and tax_slv_ssu_138.1.txt.gz available through the SILVA archive (https://www.arb-silva.de/download/archive/).

## Notes

### Competing Interest Statement

The authors have declared no competing interest.

## References

1. Díaz S, Settele J, Brondízio ES, Ngo HT, Guèze M, Agard J, et al. Summary for policymakers of the global assessment report on biodiversity and ecosystem services of the Intergovernmental Science-Policy Platform on Biodiversity and Ecosystem Services [Internet]. Bonn, Germany; 2019. Available from: https://doi.org/10.5281/zenodo.3553579

2. WWF. Living Planet Report 2020 - Bending the curve of biodiversity loss. Almond Rea, Grooten M, Petersen T, editors. Gland, Switzerland; 2020.

3. Pettorelli N, Graham NAJ, Seddon N, Maria da Cunha Bustamante M, Lowton MJ, Sutherland WJ, et al. Time to integrate global climate change and biodiversity science-policy agendas. J Appl Ecol. 2021;58:2384–93.

4. Kubiszewski I, Costanza R, Anderson S, Sutton P. The future value of ecosystem services: Global scenarios and national implications. Ecosyst Serv [Internet]. 2017;26:289–301. Available from: http://dx.doi.org/10.1016/j.ecoser.2017.05.004

5. Smith MB, Rocha AM, Smillie CS, Olesen SW, Paradis C, Wu L, et al. Natural Bacterial Communities Serve as Quantitative Geochemical Biosensors. MBio. 2015;6:e00326–15.

6. Pawlowski J, Lejzerowicz F, Apotheloz-Perret-Gentil L, Visco JA, Esling P. Protist metabarcoding and environmental biomonitoring: Time for change. Eur J Protistol [Internet]. 2016;55:12–25. Available from: http://dx.doi.org/10.1016/j.ejop.2016.02.003

7. Cordier T, Lanzén A, Apothéloz-Perret-Gentil L, Stoeck T, Pawlowski J. Embracing Environmental Genomics and Machine Learning for Routine Biomonitoring. Trends Microbiol [Internet]. 2019;27:387–97. Available from: https://doi.org/10.1016/j.tim.2018.10.012

8. Sagova-Mareckova M, Boenigk J, Bouchez A, Cermakova K, Chonova T, Cordier T, et al. Expanding ecological assessment by integrating microorganisms into routine freshwater biomonitoring. Water Res [Internet]. 2021;191:116767. Available from: https://doi.org/10.1016/j.watres.2020.116767

9. Stat M, Huggett MJ, Bernasconi R, Dibattista JD, Berry TE, Newman SJ, et al. Ecosystem biomonitoring with eDNA: Metabarcoding across the tree of life in a tropical marine environment. Sci Rep [Internet]. 2017;7:1–11. Available from: http://dx.doi.org/10.1038/s41598-017-12501-5

10. Meisel JS, Hannigan GD, Tyldsley AS, SanMiguel AJ, Hodkinson BP, Zheng Q, et al. Skin Microbiome Surveys Are Strongly Influenced by Experimental Design. J Invest Dermatol [Internet]. 2016;136:947–56. Available from: http://dx.doi.org/10.1016/j.jid.2016.01.016

11. Lozupone CA, Stombaugh J, Gonzalez A, Ackermann G, Wendel D, Vázquez-Baeza Y, et al. Meta-analyses of studies of the human microbiota. Genome Res. 2013;23:1704–14.

12. Pinto AJ, Raskin L. PCR biases distort bacterial and archaeal community structure in pyrosequencing datasets. PLoS One. 2012;7.

13. Laursen MF, Dalgaard MD, Bahl MI. Genomic GC-content affects the accuracy of 16S rRNA gene sequencing bsed microbial profiling due to PCR bias. Front Microbiol. 2017;8:1–8.

14. Walker AW, Martin JC, Scott P, Parkhill J, Flint HJ, Scott KP. 16S rRNA gene-based profiling of the human infant gut microbiota is strongly influenced by sample processing and PCR primer choice. Microbiome [Internet]. 2015;3:1–11. Available from: http://dx.doi.org/10.1186/s40168-015-0087-4

15. Wooley JC, Godzik A, Friedberg I. A primer on metagenomics. PLoS Comput Biol. 2010;6:e1000667.

16. Almeida OGG, De Martinis ECP. Bioinformatics tools to assess metagenomic data for applied microbiology. Appl Microbiol Biotechnol. 2019;103:69–82.

17. Bashiardes S, Zilberman-Schapira G, Elinav E. Use of metatranscriptomics in microbiome research. Bioinform Biol Insights. 2016;10:19–25.

18. Shakya M, Lo CC, Chain PSG. Advances and challenges in metatranscriptomic analysis. Front Genet. 2019;10:1–10.

19. Yilmaz P, Kottmann R, Pruesse E, Quast C, Glöckner FO. Analysis of 23S rRNA genes in metagenomes - A case study from the Global Ocean Sampling Expedition. Syst Appl Microbiol [Internet]. 2011;34:462–9. Available from: http://dx.doi.org/10.1016/j.syapm.2011.04.005

20. Tessler M, Neumann JS, Afshinnekoo E, Pineda M, Hersch R, Velho LFM, et al. Large-scale differences in microbial biodiversity discovery between 16S amplicon and shotgun sequencing. Sci Rep. 2017;7:1–14.

21. Shakya M, Quince C, Campbell JH, Yang ZK, Schadt CW, Podar M. Comparative metagenomic and rRNA microbial diversity characterization using archaeal and bacterial synthetic communities. Environ Microbiol. 2013;15:1882–99.

22. Logares R, Sunagawa S, Salazar G, Cornejo-Castillo FM, Ferrera I, Sarmento H, et al. Metagenomic 16S rDNA Illumina tags are a powerful alternative to amplicon sequencing to explore diversity and structure of microbial communities. Environ Microbiol. 2014;16:2659–71.

23. Shah N, Tang H, Doak TG, Ye Y. Comparing bacterial communities inferred from 16S rRNA gene sequencing and shotgun metagenomics. Pacific Symp Biocomput 2011. 2010;165–76.

24. Brumfield KD, Huq A, Colwell RR, Olds JL, Leddy MB. Microbial resolution of whole genome shotgun and 16S amplicon metagenomic sequencing using publicly available NEON data. PLoS One. 2020;15:1–21.

25. Li F, Henderson G, Sun X, Cox F, Janssen PH, Guan LL. Taxonomic assessment of rumen microbiota using total RNA and targeted amplicon sequencing approaches. Front Microbiol. 2016;7.

26. Bang-Andreasen T, Anwar MZ, Lanzén A, Kjøller R, Rønn R, Ekelund F, et al. Total RNA sequencing reveals multilevel microbial community changes and functional responses to wood ash application in agricultural and forest soil. FEMS Microbiol Ecol. 2020;96:1–13.

27. Li F, Guan LL. Metatranscriptomic profiling reveals linkages between the active rumen microbiome and feed efficiency in beef cattle. Appl Environ Microbiol. 2017;83:1–16.

28. Urich T, Lanzén A, Qi J, Huson DH, Schleper C, Schuster SC. Simultaneous assessment of soil microbial community structure and function through analysis of the meta-transcriptome. PLoS One. 2008;3:e2527.

29. Turner TR, Ramakrishnan K, Walshaw J, Heavens D, Alston M, Swarbreck D, et al. Comparative metatranscriptomics reveals kingdom level changes in the rhizosphere microbiome of plants. ISME J. 2013;7:2248–58.

30. Xue Y, Lanzén A, Jonassen I. Reconstructing ribosomal genes from large scale total RNA meta-transcriptomic data. Bioinformatics. 2020;36:3365–71.

31. Hempel CA, Wright N, Harvie J, Hleap JS, Adamowicz SJ, Steinke D. Metagenomics versus total RNA sequencing: most accurate data-processing tools, microbial identification accuracy, and perspectives for freshwater assessments. Nucleic Acids Res [Internet]. 2022; Available from: https://doi.org/10.1093/nar/gkac689

32. Uyaguari-Diaz MI, Chan M, Chaban BL, Croxen MA, Finke JF, Hill JE, et al. A comprehensive method for amplicon-based and metagenomic characterization of viruses, bacteria, and eukaryotes in freshwater samples. Microbiome [Internet]. 2016;4:1–19. Available from: http://dx.doi.org/10.1186/s40168-016-0166-1

33. Cordier T, Alonso-Sáez L, Apothéloz-Perret-Gentil L, Aylagas E, Bohan DA, Bouchez A, et al. Ecosystems monitoring powered by environmental genomics: A review of current strategies with an implementation roadmap. Mol Ecol. 2021;30:2937–58.

34. Leese F, Bouchez A, Abarenkov K, Altermatt F, Borja Á, Bruce K, et al. Why We Need Sustainable Networks Bridging Countries, Disciplines, Cultures and Generations for Aquatic Biomonitoring 2.0: A Perspective Derived From the DNAqua-Net COST Action. Adv Ecol Res. 2018;58:63–99.

35. Witten IH, Frank E. Data Mining: Practical Machine Learning Tools and Techniques. 2nd ed. San Francisco: Elsevier Inc.; 2005.

36. Greener JG, Kandathil SM, Moffat L, Jones DT. A guide to machine learning for biologists. Nat Rev Mol Cell Biol [Internet]. 2022;23:40–55. Available from: http://dx.doi.org/10.1038/s41580-021-00407-0

37. Ghannam RB, Techtmann SM. Machine learning applications in microbial ecology, human microbiome studies, and environmental monitoring. Comput Struct Biotechnol J [Internet]. 2021;19:1092–107. Available from: https://doi.org/10.1016/j.csbj.2021.01.028

38. Crisci C, Ghattas B, Perera G. A review of supervised machine learning algorithms and their applications to ecological data. Ecol Modell [Internet]. 2012;240:113–22. Available from: http://dx.doi.org/10.1016/j.ecolmodel.2012.03.001

39. Good SP, Urycki DR, Crump BC. Predicting Hydrologic Function With Aquatic Gene Fragments. Water Resour Res. 2018;54:2424–35.

40. Cordier T, Forster D, Dufresne Y, Martins CIM, Stoeck T, Pawlowski J. Supervised machine learning outperforms taxonomy-based environmental DNA metabarcoding applied to biomonitoring. Mol Ecol Resour. 2018;18:1381–91.

41. Glasl B, Bourne DG, Frade PR, Thomas T, Schaffelke B, Webster NS. Microbial indicators of environmental perturbations in coral reef ecosystems. Microbiome. 2019;7:1–13.

42. Cordier T, Esling P, Lejzerowicz F, Visco JA, Ouadahi A, Martins C, et al. Predicting the Ecological Quality Status of Marine Environments from eDNA Metabarcoding Data Using Supervised Machine Learning. Environ Sci Technol. 2017;51:9118–26.

43. Frühe L, Cordier T, Dully V, Breiner H-W, Lentendu G, Pawlowski J, et al. Supervised machine learning is superior to indicator value inference in monitoring the environmental impacts of salmon aquaculture using eDNA metabarcodes. Mol Ecol. 2020;

44. Dully V, Balliet H, Frühe L, Däumer M, Thielen A, Gallie S, et al. Robustness, sensitivity and reproducibility of eDNA metabarcoding as an environmental biomonitoring tool in coastal salmon aquaculture – An inter-laboratory study. Ecol Indic. 2021;121.

45. Gerhard WA, Gunsch CK. Metabarcoding and machine learning analysis of environmental DNA in ballast water arriving to hub ports. Environ Int [Internet]. 2019;124:312– Available from: https://doi.org/10.1016/j.envint.2018.12.038

46. Hermans SM, Buckley HL, Case BS, Curran-Cournane F, Taylor M, Lear G. Using soil bacterial communities to predict physico-chemical variables and soil quality. Microbiome. 2020;8:1–13.

47. Chang HX, Haudenshield JS, Bowen CR, Hartman GL. Metagenome-wide association study and machine learning prediction of bulk soil microbiome and crop productivity. Front Microbiol. 2017;8:1–11.

48. Piggott JJ, Salis RK, Lear G, Townsend CR, Matthaei CD. Climate warming and agricultural stressors interact to determine stream periphyton community composition. Glob Chang Biol. 2015;21:206–22.

49. Mack L, Buchner D, Brasseur M V., Leese F, Piggott JJ, Tiegs SD, et al. Fine sediment and the insecticide chlorantraniliprole inhibit organic matter decomposition in streams through different pathways. Freshw Biol. 2022;

50. Haase P, Frenzel M, Klotz S, Musche M, Stoll S. The long-term ecological research (LTER) network: Relevance, current status, future perspective and examples from marine, freshwater and terrestrial long-term observation. Ecol Indic. 2016;100:1–3.

51. Mirtl M, T. Borer E, Djukic I, Forsius M, Haubold H, Hugo W, et al. Genesis, goals and achievements of Long-Term Ecological Research at the global scale: A critical review of ILTER and future directions. Sci Total Environ. 2018;626:1439–62.

52. Buchner D, Beermann AJ, Leese F, Weiss M. Cooking small and large portions of “biodiversity-soup”: Miniaturized DNA metabarcoding PCRs perform as good as large-volume PCRs. Ecol Evol. 2021;11:9092–9.

53. Zizka VMA, Elbrecht V, Macher JN, Leese F. Assessing the influence of sample tagging and library preparation on DNA metabarcoding. Mol Ecol Resour. 2019;19:893–9.

54. Caporaso JG, Lauber CL, Walters WA, Berg-Lyons D, Lozupone CA, Turnbaugh PJ, et al. Global patterns of 16S rRNA diversity at a depth of millions of sequences per sample. Proc Natl Acad Sci U S A. 2011;108:4516–22.

55. Frey B, Rime T, Phillips M, Stierli B, Hajdas I, Widmer F, et al. Microbial diversity in European alpine permafrost and active layers. FEMS Microbiol Ecol. 2016;92:1–17.

56. Buchner D, Macher T-H, Leese F. APSCALE: advanced pipeline for simple yet comprehensive analyses of DNA Meta-barcoding data. Bioinformatics. 2022;7:1–3.

57. Rognes T, Flouri T, Nichols B, Quince C, Mahé F. VSEARCH: A versatile open source tool for metagenomics. PeerJ. 2016;2016:1–22.

58. Martin M. Cutadapt removes adapter sequences from high-throughput sequencing reads. EMBnet.journal. 2011;17:10–2.

59. Frøslev TG, Kjøller R, Bruun HH, Ejrnæs R, Brunbjerg AK, Pietroni C, et al. Algorithm for post-clustering curation of DNA amplicon data yields reliable biodiversity estimates. Nat Commun [Internet]. 2017;8. Available from: http://dx.doi.org/10.1038/s41467-017-01312-x

60. McLaren MR, Callahan BJ. Silva 138.1 prokaryotic SSU taxonomic training data formatted for DADA2 [Internet]. Zenodo; 2021. Available from: https://doi.org/10.5281/zenodo.4587955

61. Abarenkov K, Zirk A, Piirmann T, Pöhönen R, Ivanov F, Nilsson RH, et al. UNITE general FASTA release for eukaryotes [Internet]. 2021. Available from: https://dx.doi.org/10.15156/BIO/1280127

62. Bolger AM, Lohse M, Usadel B. Trimmomatic: A flexible trimmer for Illumina sequence data. Bioinformatics. 2014;30:2114–20.

63. Bankevich A, Nurk S, Antipov D, Gurevich AA, Dvorkin M, Kulikov AS, et al. SPAdes: A new genome assembly algorithm and its applications to single-cell sequencing. J Comput Biol. 2012;19:455–77.

64. Li D, Liu CM, Luo R, Sadakane K, Lam TW. MEGAHIT: An ultra-fast single-node solution for large and complex metagenomics assembly via succinct de Bruijn graph. Bioinformatics. 2015;31:1674–6.

65. Li H, Durbin R. Fast and accurate short read alignment with Burrows-Wheeler transform. Bioinformatics. 2009;25:1754–60.

66. Li H, Handsaker B, Wysoker A, Fennell T, Ruan J, Homer N, et al. The Sequence Alignment/Map format and SAMtools. Bioinformatics. 2009;25:2078–9.

67. Quast C, Pruesse E, Yilmaz P, Gerken J, Schweer T, Yarza P, et al. The SILVA ribosomal RNA gene database project: Improved data processing and web-based tools. Nucleic Acids Res. 2013;41:590–6.

68. Wood DE, Lu J, Langmead B. Improved metagenomic analysis with Kraken 2. Genome Biol. Genome Biology; 2019;20:1–13.

69. Van Rossum G, Drake FL. Python 3 Reference Manual. Scotts Valley, CA: CreateSpace; 2009.

70. Reback J, jbrockmendel, McKinney W, den Bossche J Van, Augspurger T, Cloud P, et al. pandas-dev/pandas: Pandas 1.3.5 [Internet]. Zenodo; 2021. Available from: https://doi.org/10.5281/zenodo.5774815

71. Harris CR, Millman KJ, van der Walt SJ, Gommers R, Virtanen P, Cournapeau D, et al. Array programming with NumPy. Nature [Internet]. C}; 2020;585:357–62. Available from: https://doi.org/10.1038/s41586-020-2649-2

72. Gloor GB, Macklaim JM, Pawlowsky-Glahn V, Egozcue JJ. Microbiome datasets are compositional: And this is not optional. Front Microbiol. 2017;8:1–6.

73. The scikit-bio development team. scikit-bio: A Bioinformatics Library for Data Scientists, Students, and Developers [Internet]. 2020. Available from: http://scikit-bio.org

74. Pedregosa F, Varoquaux G, Gramfort A, Michel V, Thirion B, Grisel O, et al. Scikit-learn: Machine Learning in Python. J Mach Learn Res. 2011;12:2825–30.

75. Chen T, Guestrin C. XGBoost: A Scalable Tree Boosting System. Proc 22nd ACM SIGKDD Int Conf Knowl Discov Data Min. New York, NY, USA: ACM; 2016. p. 785–94.

76. Virtanen P, Gommers R, Oliphant TE, Haberland M, Reddy T, Cournapeau D, et al. SciPy 1.0: fundamental algorithms for scientific computing in Python. Nat Methods. 2020;17:261–72.

77. Obiol A, Giner CR, Sánchez P, Duarte CM, Acinas SG, Massana R. A metagenomic assessment of microbial eukaryotic diversity in the global ocean. Mol Ecol Resour. 2020;20:718– 31.

78. Chan CS, Chan KG, Tay YL, Chua YH, Goh KM. Diversity of thermophiles in a Malaysian hot spring determined using 16S rRNA and shotgun metagenome sequencing. Front Microbiol. 2015;6:1–15.

79. Laudadio I, Fulci V, Palone F, Stronati L, Cucchiara S, Carissimi C. Quantitative Assessment of Shotgun Metagenomics and 16S rDNA Amplicon Sequencing in the Study of Human Gut Microbiome. Omi A J Integr Biol. 2018;22:248–54.

80. Yan YW, Jiang QY, Wang JG, Zhu T, Zou B, Qiu QF, et al. Microbial communities and diversities in mudflat sediments analyzed using a modified metatranscriptomic method. Front Microbiol. 2018;9:1–15.

81. Quince C, Walker AW, Simpson JT, Loman NJ, Segata N. Shotgun metagenomics, from sampling to analysis. Nat Biotechnol. 2017;35:833–44.

82. Hempel CA, Carson SEE, Elliott TA, Adamowicz SJ. Reconstruction of Small Subunit Ribosomal RNA from High-Throughput Sequencing Data : A Comparative Study of Metagenomics and Total RNA Sequencing. bioRxiv. 2022;1–31.

83. Marcos-Zambrano LJ, Karaduzovic-Hadziabdic K, Loncar Turukalo T, Przymus P, Trajkovik V, Aasmets O, et al. Applications of Machine Learning in Human Microbiome Studies: A Review on Feature Selection, Biomarker Identification, Disease Prediction and Treatment. Front Microbiol. 2021;12.

84. Douglas GM, Hansen R, Jones CMA, Dunn KA, Comeau AM, Bielawski JP, et al. Multi-omics differentially classify disease state and treatment outcome in pediatric Crohn ‘ s disease. Microbiome; 2018;1–12.

85. Ai D, Pan H, Han R, Li X, Liu G, Xia LC. Using decision tree aggregation with random forest model to identify gut microbes associated with colorectal cancer. Genes (Basel). 2019;10.

86. Wu H, Cai L, Li D, Wang X, Zhao S, Zou F, et al. Metagenomics Biomarkers Selected for Prediction of Three Different Diseases in Chinese Population. Biomed Res Int. 2018;2018.

87. Hacilar H, Nalbantoglu OU, Bakir-Gungor B. Machine Learning Analysis of Inflammatory Bowel Disease-Associated Metagenomics Dataset. UBMK 2018 - 3rd Int Conf Comput Sci Eng. 2018;434–8.

88. Knight R, Vrbanac A, Taylor BC, Aksenov A, Callewaert C, Debelius J, et al. Best practices for analysing microbiomes. Nat Rev Microbiol [Internet]. 2018;16:410–22. Available from: http://dx.doi.org/10.1038/s41579-018-0029-9

89. Callahan BJ, McMurdie PJ, Holmes SP. Exact sequence variants should replace operational taxonomic units in marker-gene data analysis. ISME J [Internet]. Nature Publishing Group; 2017;11:2639–43. Available from: http://dx.doi.org/10.1038/ismej.2017.119

90. Glassman SI, Martiny JBH. Broadscale Ecological Patterns Are Robust to Use of Exact. mSphere. 2018;3:e00148–18.

91. Vera-Gargallo B, Chowdhury TR, Brown J, Fansler SJ, Durán-Viseras A, Sánchez-Porro C, et al. Spatial distribution of prokaryotic communities in hypersaline soils. Sci Rep. 2019;9:1– 12.

92. Kang W, Anslan S, Börner N, Schwarz A, Schmidt R, Künzel S, et al. Diatom metabarcoding and microscopic analyses from sediment samples at Lake Nam Co, Tibet: The effect of sample-size and bioinformatics on the identified communities. Ecol Indic. 2021;121.

93. Tapolczai K, Keck F, Bouchez A, Rimet F, Kahlert M, Vasselon V. Diatom DNA Metabarcoding for Biomonitoring: Strategies to Avoid Major Taxonomical and Bioinformatical Biases Limiting Molecular Indices Capacities. Front Ecol Evol. 2019;7:1–15.

94. Joos L, Beirinckx S, Haegeman A, Debode J, Vandecasteele B, Baeyen S, et al. Daring to be differential: metabarcoding analysis of soil and plant-related microbial communities using amplicon sequence variants and operational taxonomical units. BMC Genomics. 2020;21:1–17.

95. Caruso V, Song X, Asquith M, Karstens L. Performance of Microbiome Sequence Inference Methods in Environments with Varying Biomass. mSystems. 2019;4.

96. Roy J, Mazel F, Sosa-Hernández MA, Dueñas JF, Hempel S, Zinger L, et al. The relative importance of ecological drivers of arbuscular mycorrhizal fungal distribution varies with taxon phylogenetic resolution. New Phytol. 2019;224:936–48.

97. Tedersoo L, Bahram M, Zinger L, Nilsson RH, Kennedy PG, Yang T, et al. Best practices in metabarcoding of fungi: From experimental design to results. Mol Ecol. 2022;31:2769–95.

98. Philippot L, Andersson SGE, Battin TJ, Prosser JI, Schimel JP, Whitman WB, et al. The ecological coherence of high bacterial taxonomic ranks. Nat Rev Microbiol [Internet]. 2010;8:523–9. Available from: http://dx.doi.org/10.1038/nrmicro2367

99. Choe YH, Kim M, Lee YK. Distinct Microbial Communities in Adjacent Rock and Soil Substrates on a High Arctic Polar Desert. Front Microbiol. 2021;11:1–15.

100. Auladell A, Barberán A, Logares R, Garcés E, Gasol JM, Ferrera I. Seasonal niche differentiation among closely related marine bacteria. ISME J. 2022;16:178–89.

101. Dillies MA, Rau A, Aubert J, Hennequet-Antier C, Jeanmougin M, Servant N, et al. A comprehensive evaluation of normalization methods for Illumina high-throughput RNA sequencing data analysis. Brief Bioinform. 2013;14:671–83.

102. Pereira MB, Wallroth M, Jonsson V, Kristiansson E. Comparison of normalization methods for the analysis of metagenomic gene abundance data. BMC Genomics. 2018;19:1–17.

103. Weiss S, Xu ZZ, Peddada S, Amir A, Bittinger K, Gonzalez A, et al. Normalization and microbial differential abundance strategies depend upon data characteristics. Microbiome [Internet]. 2017;5:27. Available from: http://www.ncbi.nlm.nih.gov/pubmed/28253908%0Ahttp://www.pubmedcentral.nih.gov/articlerender.fcgi?artid=PMC5335496

104. Koh H, Li Y, Zhan X, Chen J, Zhao N. A distance-based kernel association test based on the generalized linear mixed model for correlated microbiome studies. Front Genet. 2019;10:1–14.

105. Buchner D, Beermann AJ, Laini A, Rolauffs P, Vitecek S, Hering D, et al. Analysis of 13,312 benthic invertebrate samples from German streams reveals minor deviations in ecological status class between abundance and presence/absence data. PLoS One. 2019;14:1–18.

106. Farinella R, Rizzato C, Bottai D, Bedini A, Gemignani F, Landi S, et al. Maternal anthropometric variables and clinical factors shape neonatal microbiome. Sci Rep [Internet]. 2022;12:1–10. Available from: https://doi.org/10.1038/s41598-022-06792-6

107. Muletz Wolz CR, Yarwood SA, Campbell Grant EH, Fleischer RC, Lips KR. Effects of host species and environment on the skin microbiome of Plethodontid salamanders. J Anim Ecol. 2018;87:341–53.

108. Knowles SCL, Eccles RM, Baltrūnaité L. Species identity dominates over environment in shaping the microbiota of small mammals. Ecol Lett. 2019;22:826–37.

109. Tavalire HF, Christie DM, Leve LD, Ting N, Cresko WA, Bohannan BJM. Shared environment and genetics shape the gut microbiome after infant adoption. MBio. 2021;12.

110. Kask O, Kyman S, Conn KA, Gormley J, Gardner J, Johns RA, et al. Environmental Exposures Influence Nasal Microbiome Composition in a Longitudinal Study of Division I Collegiate Athletes. bioRxiv. 2020;

111. Khachatryan L, de Leeuw RH, Kraakman MEM, Pappas N, te Raa M, Mei H, et al. Taxonomic classification and abundance estimation using 16S and WGS—A comparison using controlled reference samples. Forensic Sci Int Genet [Internet]. 2020;46:102257. Available from: https://doi.org/10.1016/j.fsigen.2020.102257

112. Oudah M, Henschel A. Taxonomy-aware feature engineering for microbiome classification. BMC Bioinformatics. 2018;19:1–13.

113. Lanzén A, Mendibil I, Borja A, Laura Alonse Saez. A microbial mandala for environmental monitoring – predicting multiple impacts on estuarine prokaryote communities of the Bay of Biscay. Mol Ecol. 2020;

